# “Replication stalling activates SSB for recruitment of DNA damage tolerance factors”

**DOI:** 10.1101/2022.05.24.493288

**Authors:** Elizabeth S. Thrall, Sadie C. Piatt, Seungwoo Chang, Joseph J. Loparo

**Author notes:** Equal contribution (Elizabeth S. Thrall, Sadie C. Piatt). Department of Chemistry, Fordham University, Bronx, NY. **Author Contributions:** E.S.T., S.C.P., and J.J.L. designed research; E.S.T. and S.C.P. performed research; E.S.T., S.C.P., and S.C. contributed new reagents; E.S.T., S.C.P., and J.J.L. analyzed data; and E.S.T., S.C.P., S.C., and J.J.L. wrote the paper. **Competing Interest Statement:** The authors declare no competing interests.

## Abstract

Translesion synthesis (TLS) polymerases bypass DNA lesions that block replicative polymerases, allowing cells to tolerate DNA damage encountered during replication. It is well known that most bacterial TLS polymerases must interact with the sliding clamp processivity factor to carry out TLS, but recent work in *Escherichia coli* has revealed that single-stranded DNA-binding protein (SSB) plays a key role in enriching the TLS polymerase Pol IV at stalled replication forks in the presence of DNA damage. It remains unclear how this interaction with SSB enriches Pol IV in a stalling-dependent manner given that SSB is always present at the replication fork. In this study we use single-molecule imaging in live *E. coli* cells to investigate this SSB-dependent enrichment of Pol IV. We find that Pol IV is enriched through its interaction with SSB in response to a range of different replication stresses, and that changes in SSB dynamics at stalled forks may explain this conditional Pol IV enrichment. Finally, we show that other SSB-interacting proteins (SIPs) are likewise selectively enriched in response to replication perturbations, suggesting that this mechanism is likely a general one for enrichment of repair factors near stalled replication forks.

## Introduction

Translesion synthesis (TLS) is a DNA damage tolerance pathway, conserved across all domains of life, that can rescue replication forks stalled at sites of DNA damage.(1, 2) In this process, a TLS polymerase exchanges with the replicative polymerase and continues DNA synthesis past the lesion on the templating strand. By allowing processive replication to continue downstream of the lesion, TLS helps cells avoid the deleterious consequences of a stalled replication fork, including replication fork collapse and toxic double-strand DNA breaks. Proper regulation of TLS is critical, however, because most TLS polymerases are error-prone and can introduce harmful mutations if their access to the DNA template is not restricted.(3)

Previously we used single-molecule live cell imaging to gain insight into the regulation of the most abundant TLS polymerase in *Escherichia coli*, Pol IV.(4) We found that Pol IV is only modestly enriched near sites of replication under normal growth conditions but becomes strongly colocalized with the replisome when cells are treated with the DNA damaging agent methyl methanesulfonate (MMS). This lack of enrichment of Pol IV near replication forks in the absence of DNA damage likely limits the access of Pol IV to the DNA template during normal replication in order to minimize mutagenesis.

It is well established that Pol IV and other TLS polymerases interact with the replication processivity factor, or the sliding clamp, and that this interaction is critical for TLS.(5, 6) In addition to TLS polymerases, clamps bind a number of other proteins involved in DNA replication and repair.(7–9) The *E. coli* β clamp is a dimer with two equivalent binding cleft sites for clamp interacting proteins.(10) The replicative polymerase, Pol III, binds to both sites through its α and ε subunits,(11) which can occlude Pol IV access.(12) Less is known, however, about the importance of other protein-protein interactions in TLS. Previous studies have reported that Pol IV also interacts with single-stranded DNA-binding protein (SSB)(13) and both the RecA recombinase and the UmuD subunit of the TLS polymerase Pol V.(14, 15) Like the clamp, *E. coli* SSB is known to interact with a wide range of DNA replication and repair proteins, which bind to the conserved C-terminal peptide.(16)

Recently our lab showed that the Pol IV-SSB interaction enriches Pol IV near replication forks stalled by DNA damage.(17) Locally concentrating Pol IV enables it to outcompete Pol III and gain access to the β clamp, which is critical for TLS on both the leading and lagging strands. The interaction with SSB might also help Pol IV outcompete other clamp-binding proteins that do not interact with SSB. The clustering of the SSB C-terminal tail appears to be a key feature required for Pol IV localization, likely due to the relatively weak (low micromolar) binding affinity between Pol IV and a single SSB C-terminal tail.

The role of the SSB interaction in enriching Pol IV near replication forks raises several new questions. First, we initially observed this enrichment when cells were treated with MMS, which generates DNA lesions that Pol IV is capable of bypassing.(4) Is Pol IV only enriched near replication forks in the presence of such “cognate” DNA lesions, or is enrichment a general consequence of replication stalling independent of the cause? Second, SSB binds transiently exposed ssDNA on the lagging strand during replication; thus, it is a constitutive component of normal replication forks. Yet we observed little Pol IV enrichment near replication forks in the absence of DNA damage.(4) What features differentiate SSB at stalled forks from SSB at moving forks such that Pol IV is selectively enriched upon stalling? Finally, *E. coli* SSB interacts with at least 17 different SSB interacting proteins (SIPs).(16, 18) Are other SIPs selectively enriched near stalled replication forks upon replication stalling, or is this mechanism unique to Pol IV?

In this study we use single-molecule fluorescence imaging in live *E. coli* cells to address these questions. We find that Pol IV is enriched near replication forks in the presence of “non-cognate” forms of DNA damage, meaning lesions that it cannot bypass, as well as upon nucleotide depletion, suggesting that Pol IV enrichment is a general consequence of replication stalling. Further, we show that this enrichment requires interactions with the β clamp and SSB, with SSB playing the major role. By imaging single SSB molecules, we find that SSB dynamics change upon replication stress, with an increase in the relative population of statically bound SSB molecules and a concomitant decrease in the mobile population. In addition to a modest increase in SSB copy number at replication forks, we observe a substantial increase in the SSB binding lifetime upon replication perturbation, providing a possible mechanism for selective Pol IV enrichment. Finally, we show that two other SIPs, PriA and RecG, are also enriched near sites of replication only upon DNA damage, suggesting that changes in SSB dynamics may provide a general mechanism for enriching factors that can respond to stalled replication forks.

## Results

### Pol IV is strongly enriched at replication forks in the presence of cognate DNA damage

Previously we constructed and validated a functional C-terminal fusion of Pol IV to the photoactivatable fluorescent protein PAmCherry(19) at the endogenous *dinB* locus, where *dinB* is the gene encoding Pol IV.(4) To prevent large changes in copy number in response to DNA damage, we introduced this fusion in the *lexA51* strain background, in which the SOS DNA damage response is constitutively activated due to a truncation of the LexA repressor(20, 21); the SOS response increases Pol IV levels by approximately ten-fold.(5, 22) To visualize sites of DNA replication in the cell, we introduced an orthogonal C-terminal fusion of single-stranded DNA-binding protein (SSB) to the yellow fluorescent protein variant mYPet.(23) Because replacement of all SSB copies with a C-terminal fusion has been reported to be lethal(24) or to impair protein-protein interactions,(18) we introduced this SSB fusion as a second copy at the IPTG-inducible *lacZ* locus.

To determine whether and under what conditions Pol IV is colocalized with sites of replication, we imaged Pol IV-PAmCherry in live *E. coli* cells in a custom-built single-molecule fluorescence microscope using particle-tracking photoactivation localization microscopy (PALM).(25, 26) In brief, a 405 nm near-UV laser was used to photoactivate PAmCherry molecules from a dark (non-fluorescent) state to a bright (fluorescent) state that was excited using 561 nm laser excitation. By adjusting power density at 405 nm, the photoactivation rate was controlled such that at most one PAmCherry molecule was photoactivated at a time (Figures 1A and 1C). The motion of each individual Pol IV-PAmCherry molecule was tracked until the molecule photobleached, converting it irreversibly to a dark state. In the same cells, we used 514 nm laser excitation to excite SSB-mYPet, which formed foci at replication forks (Figures 1B and 1D) as previously reported.(27) Previously we observed two populations of Pol IV in normally-growing *E. coli* cells, a fast-moving diffusive population and an immobile statically-bound population.(4) This immobile population should include molecules specifically bound to the β clamp or to SSB, although it may also include molecules that are transiently bound to DNA in a non-specific manner.(28) To resolve immobile molecules selectively, we used a long exposure time of 250 ms, which blurs out fast-moving molecules, and identified static molecules based on the width of their point spread function (PSF).(4, 26)

**Fig. 1.**
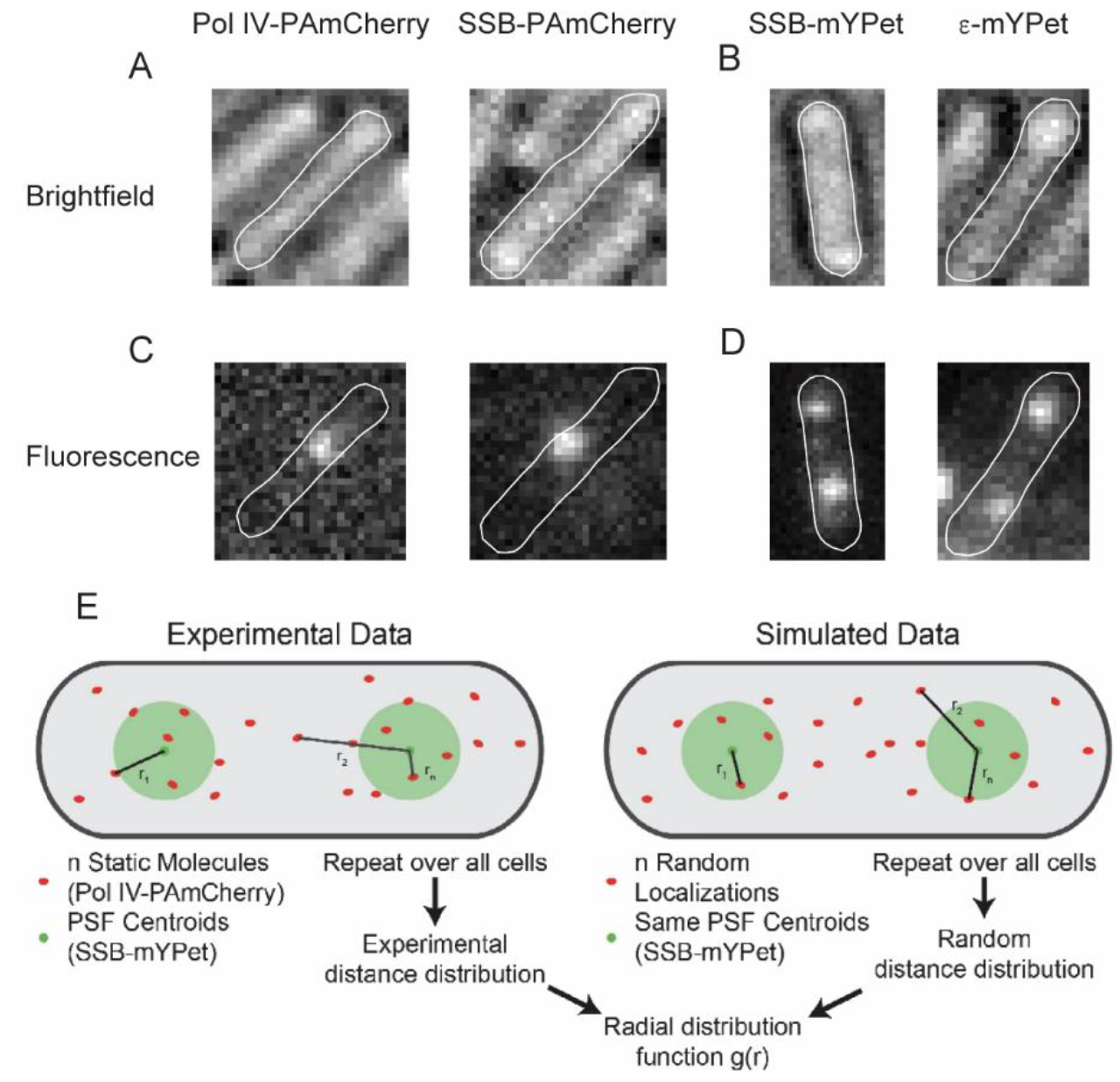
Particle-tracking PALM imaging of Pol IV-PAmCherry and SSB-PAmCherry molecules and standard near-TIRF fluorescence imaging of SSB-mYPet and Pol III ε-mYPet foci. Representative brightfield micrographs of cells containing (*A*) PAmCherry (*pamcherry1*) fusions to the endogenous copy of the Pol IV gene *dinB* (left) or a second copy of the single-stranded DNA-binding protein gene *ssb* at the *lacZ* locus (right) and (*B*) mYPet (*mypet*) fusions to a second copy of *ssb* at the *lacZ* locus (left) or the endogenous copy of the ε subunit of Pol III *dnaQ* (right). Representative fluorescence micrographs of (*C*) single activated Pol IV-PAmCherry (left) or SSB-PAmCherry (right) molecules and (*D*) SSB-mYPet (left) and ε-mYPet (right) foci recorded with 250 ms integration times with overlays of the cell outlines. (*E*) Schematic of radial distribution function analysis. Experimental and simulated random distributions of the distance between the nearest mYPet centroid and each static PAmCherry trajectory are determined. The experimental distribution is normalized by the simulated random distribution to obtain the radial distribution function *g*(*r*). (Adapted from (17).)

The degree of colocalization between these static Pol IV molecules and sites of DNA replication can be measured by calculating the distance between each Pol IV molecule and the nearest SSB focus (Figure 1E). This raw Pol IV-SSB distance distribution, however, is sensitive to the size of the cell and the number of replication forks, which affect the probability that a Pol IV molecule would appear to colocalize with an SSB focus by chance. To correct for these effects, we used radial distribution function analysis.(4, 29, 30) In this approach (see Methods for more detail), a random Pol IV-SSB distance distribution was generated by randomly simulating Pol IV localizations in each cell using the same cell outlines and the same SSB focus positions (Figure 1E). The experimentally observed distance distribution was then normalized by the random distribution, giving the radial distribution function, *g*(*r*). The magnitude of *g*(*r*) at each value of *r* reflects the fold enrichment of Pol IV at that distance from SSB relative to the random distribution. Thus *g*(*r*) values of one indicate no enrichment relative to random chance, whereas values greater than one indicate enrichment.

Consistent with our previous report, we observed little enrichment of Pol IV near replication forks in untreated *E. coli* cells (Figure 2A).(4) However, Pol IV was strongly enriched at sites of replication, with *g*(*r*) ≈ 8 at short Pol IV-SSB distances (Figure 2B), in cells treated with a 100 mM concentration of the alkylating agent methyl methanesulfonate (MMS) (see Methods). Pol IV can efficiently bypass the types of DNA damage generated by MMS, particularly the replication-blocking *N*^3^-methyladenine lesion.(17, 31) Mutating the two clamp-binding sites on Pol IV (Pol IV^R,C^), which contact the rim and cleft of the β clamp, led to a reduction in the maximum *g*(*r*) value to approximately 5 (Figure 2B).(4) Likewise, the T120P mutation, which abrogates Pol IV binding to SSB (Pol IV^T120P^), produced a larger reduction in the maximum *g*(*r*) to approximately 3 (Figure 2B).(17) From these results, we previously concluded that SSB plays the primary role in enrichment of Pol IV at stalled replication forks, whereas the interaction with the β clamp stabilizes Pol IV at the primer-template junction and is essential for synthesis past the lesion. Neither mutant was enriched near sites of replication in untreated cells, with a maximum *g*(*r*) ≈ 1 for both (Figure 2A).

**Fig. 2.**
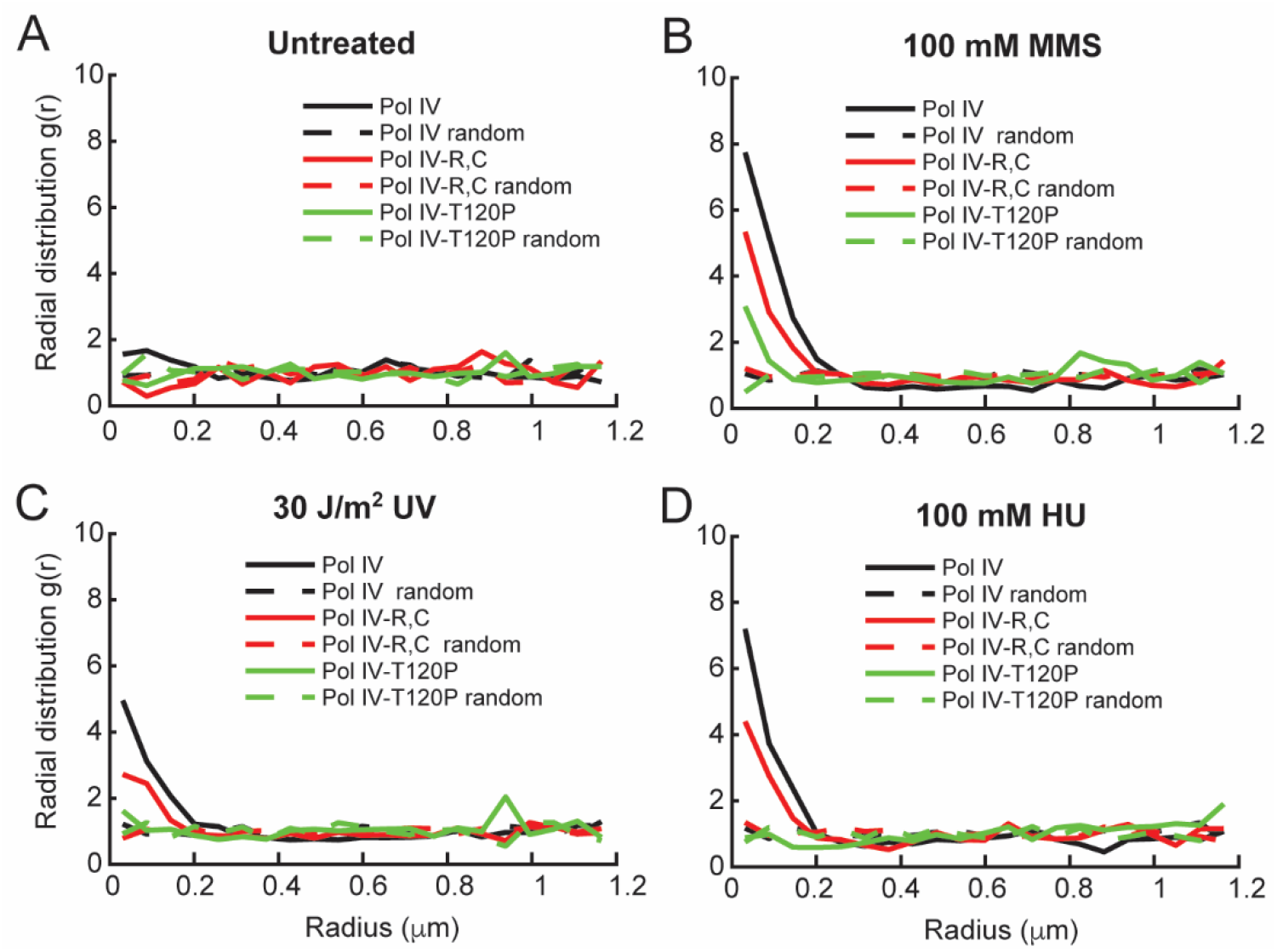
Effect of MMS, UV, and HU treatment on single-cell colocalization of Pol IV-PAmCherry and SSB-mYPet. Radial distribution function *g*(*r*) between each static Pol IV-PAmCherry track and the nearest SSB-mYPet focus for Pol IV^WT^ (black), Pol IV^R,C^ (red), and Pol IV^T120P^ (green) in (*A*) untreated cells (*N* = 1,482, 1,090, and 878 respectively) and cells treated with (*B*) 100 mM MMS (*N* = 2,471, 2,894, and 2,331 respectively), (*C*) 30 J/m^2^ UV light (*N* = 2,681, 3,261, and 1,142 respectively), and (*D*) 100 mM HU (*N* = 1,539, 1,479, and 1,207 respectively).

### Pol IV is enriched at replication forks even in the presence of non-cognate DNA damage

In contrast to MMS, there are some DNA damaging agents that generate lesions that Pol IV cannot bypass as efficiently. Next, we asked whether Pol IV would be enriched at replication forks in the presence of such non-cognate DNA lesions or whether localization requires Pol IV recognition of a cognate lesion. We treated cells with 30 J/m^2^ of 254 nm UV light, which generates strongly blocking DNA lesions including cyclobutane pyrimidine dimers and 6-4 photoproducts.(32) A similar dosage was estimated to generate approximately one cyclobutane–pyrimidine dimer per 9 kb of DNA and found to inhibit DNA replication without having a major impact on survival of WT *E. coli* cells.(33–35) We observed moderate enrichment of Pol IV at sites of replication upon UV treatment (maximum *g*(*r*) ≈ 5) (Figure 2C), consistent with a prior report.(22) Importantly, this enrichment was dependent on the same protein-protein interactions as in MMS-treated cells. The maximum *g*(*r*) value was reduced by nearly 2-fold (*g*(*r*) ≈ 3) for the Pol IV^R,C^ mutant and by approximately 2.5-fold for the Pol IV^T120P^ mutant (*g*(*r*) ≈ 2) (Figure 2C). Although the maximum value of *g*(*r*) for a given treatment condition depends on factors including the lesion density and the dynamics of Pol IV binding, these results indicate that Pol IV is enriched at replication forks through interactions with the β clamp and SSB even when it is unable to carry out TLS. They also support our view that the static population of Pol IV includes both molecules actively synthesizing DNA as well as molecules that are bound at the replication fork but not carrying out TLS.

### Pol IV is enriched at replication forks upon DNA damage-independent replication stalling

Since both cognate and non-cognate DNA damage enrich Pol IV at the replication fork in an SSB-dependent manner, our results suggest that any obstacles to processive replication may act as a molecular signal for the recruitment of Pol IV. To test this possibility, we treated cells with the ribonucleotide reductase (RNR) inhibitor hydroxyurea (HU).(36) RNR inhibition by HU depletes cellular dNTP pools, ultimately leading to replication stalling and cell death.(37) Cells lacking Pol IV are not sensitized to HU relative to WT cells (Figure S1B), indicating that Pol IV does not contribute to survival upon HU treatment.

We found that Pol IV was strongly enriched at replication forks when cells were treated with 100 mM HU (maximum *g*(*r*) ≈ 7) (Figure 2D), with a slightly lower level of enrichment upon treatment with 20 mM HU (Figure S2). To test whether Pol IV enrichment was specific to HU treatment or resulted generally from nucleotide depletion, we tested a different RNR inhibitor, guanazole.(36, 38) Pol IV enrichment was comparable for cells treated with a 100 mM concentration of guanazole and with the same concentration of HU (Figure S2), suggesting that the enrichment is a general result of nucleotide depletion and not specific to HU. Next, we asked whether the same molecular interactions were required for Pol IV enrichment upon this DNA damage-independent stalling. We measured a reduction of almost two-fold in the maximum *g*(*r*) value for the ß-binding deficient Pol IV^R,C^ mutant in comparison to Pol IV^WT^ in cells treated with 100 mM HU, comparable to the loss of enrichment observed for this mutant relative to Pol IV^WT^ upon MMS treatment (*g*(*r*) ≈ 4) (Figure 2D). Further, the SSB-binding deficient Pol IV^T120P^ mutant was not enriched at all relative to random colocalization upon HU treatment (Figure 2D). These results support a model where the Pol IV-SSB interaction plays the dominant role in Pol IV recruitment upon general replication stalling and not just in response to DNA damage-induced stalling.

### SSB dynamics change upon perturbations to replication

Taken together, our data suggest that interactions with SSB enrich Pol IV at stalled replication forks, independent of whether Pol IV is capable of resolving the stall. SSB is present at the replication fork during normal replication where it transiently binds single-stranded DNA (ssDNA) exposed on the lagging strand. Thus, our results raise the question of why Pol IV is not strongly enriched at the replication fork through interactions with SSB during normal replication, but only upon replication stalling. We reasoned that one possible mechanism could be changes in SSB behavior upon perturbations to replication.

To probe changes in the behavior of single SSB molecules upon replication stalling, we created an SSB-PAmCherry fusion, introduced as a second copy at the *lacZ* locus in the same manner as the mYPet fusion. PALM imaging allowed us to visualize individual SSB-PAmCherry molecules as bright spots in the cell (Figure 1C). As a replisome marker, we used a previously characterized fusion of mYPet to the proofreading exonuclease subunit ε (encoded by the *dnaQ* gene) of the replicative polymerase, Pol III.(4) This ε-mYPet fusion forms distinct foci at sites of replication (Figure 1D). Cells bearing the SSB-PAmCherry fusion were not sensitized to MMS treatment relative to WT cells (Figure S1A), indicating that the SSB fusion does not appear to impair TLS or other DNA damage response pathways.

To characterize SSB mobility, we imaged untreated *E. coli* cells containing this SSB-PAmCherry fusion using a short 13.3 ms exposure time. Two broad populations of SSB molecules were observed in cells: a static fraction, with an apparent diffusion coefficient *D*\* ≈ 0.08 μm^2^/s and a mobile fraction with *D*\* ≈ 1 μm^2^/s (Figure 3A); *D*\* for the static molecules is slightly larger than 0 due to the localization precision of approximately 17 nm, which leads to some apparent motion of immobile molecules.(4, 26) As for Pol IV and other DNA-binding proteins, this static fraction comprises DNA-bound molecules, whereas the mobile fraction represents free SSB molecules. The static population of SSB molecules is expected to be disproportionately localized near sites of replication, where SSB is bound to Okazaki fragments. Consistent with this prediction, SSB molecules localized within 200 nm of the nearest ε-mYPet focus are almost entirely static (Figure 3B), whereas both static and mobile populations are represented outside that radius (Figure 3C). There are several possible explanations for the presence of static SSB molecules that do not appear to colocalize with replication forks, including the presence of ssDNA gaps on incomplete Okazaki fragments or at sites of DNA repair, SSB molecules that are transiently static but are not actually DNA-bound, or missed detection of ε-mYPet foci.

**Fig. 3.**
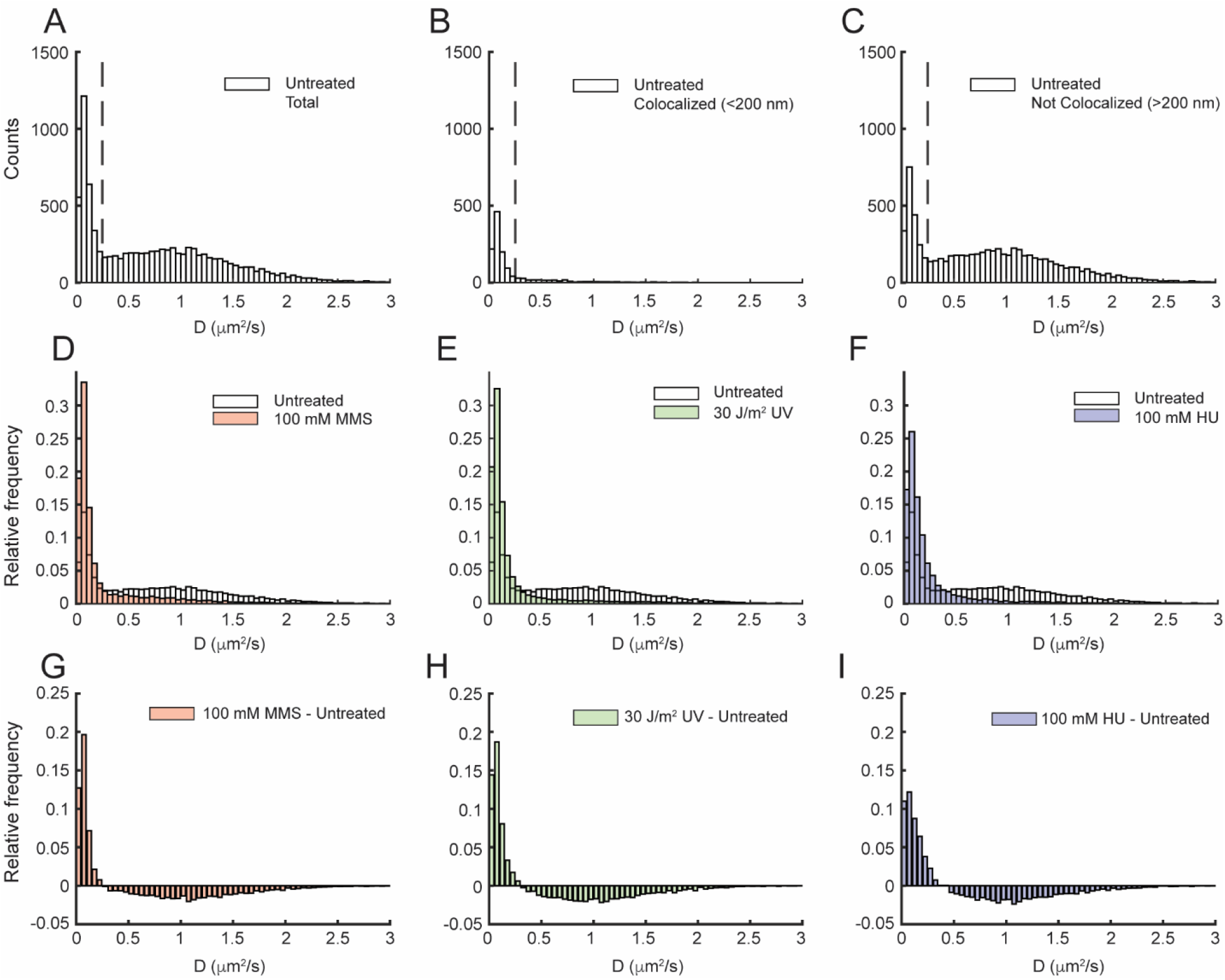
Effect of MMS, UV, and HU treatment on the apparent diffusion coefficient (*D*\*) of SSB-PAmCherry. *D*\* distributions of SSB-PAmCherry in untreated cells for (*A*) all molecules (*N* = 8,722), (*B*) molecules < 200 nm from Pol III ε-mYPet foci (*N* = 1,276), and (*C*) molecules > 200 nm from Pol III ε-mYPet foci (*N* = 7,446). The dashed lines indicate the threshold *D*\* value for bound molecules (*D*\* < 0.25 μm^2^/s). *D*\* distributions of SSB-PAmCherry for all molecules in untreated cells and cells treated with (*D*) 100 mM MMS (red, *N* = 11,797), (*E*) 30 J/m^2^ UV light (green, *N* = 16,542), and (*F*) 100 mM HU (blue, *N* = 5,690). (*G* – *I*) The difference in *D*\* distributions of SSB-PAmCherry between untreated cells and MMS-, UV-, and HU-treated cells, respectively.

Upon treatment of cells with 100 mM MMS, 30 J/m^2^ UV, or 100 mM HU, there was a dramatic reduction in the mobile fraction of SSB and a corresponding increase in the static fraction (Figure 3D – I, Tables S1 and S2). To quantify these changes, the apparent diffusion coefficient distributions were fit to a three-species model (see Figure S3 and Methods). The three populations had mean apparent diffusion coefficients in the ranges of 0.06 – 0.08 μm^2^/s, 0.15 – 0.25 μm^2^/s, and 1.0 – 1.2 μm^2^/s, with the exact values depending on the treatment condition (Table S3). The first and third populations represent stably bound and mobile SSB respectively, whereas the intermediate population may represent SSB molecules interacting transiently with DNA. In untreated cells, static molecules represented 24% of the total SSB population; this fraction doubled in cells treated with MMS, UV, or HU (to 47%, 50%, and 45% respectively). Thus, there is a dramatic change in the mobility of SSB molecules upon replication perturbation, which could act as a molecular signal to enrich Pol IV in a replication stalling-specific manner.

### The binding lifetime of SSB at the replication fork increases upon perturbations to replication

The increase in the static fraction of SSB molecules upon replication stalling suggests an increase in the average SSB copy number at the replication fork, but it may also partially reflect an increase in the average SSB binding lifetime at the replication fork. Either of these changes could contribute to a larger fraction of static SSB molecules in the distribution of apparent diffusion coefficients. To test the first of these possible changes, we carried out an analysis of SSB-mYPet foci in untreated cells and cells treated with 100 mM MMS, 30 J/m^2^ UV, and 100 mM HU. In brief, SSB-mYPet foci were fit to symmetric two-dimensional Gaussian functions and the integrated area was determined (see Methods for further details). For the 100 mM MMS and 100 mM HU conditions, increases of approximately 50% and 25% in the median integrated intensity of the SSB foci were observed. For 30 J/m^2^ UV, the increase in the median was over two-fold for the integrated SSB-mYPet intensity (Figure 4A – C, Table S4). Under all treatment conditions, there was a minimal change in the mean number of SSB-mYPet foci per cell (Table S5).

**Fig. 4.**
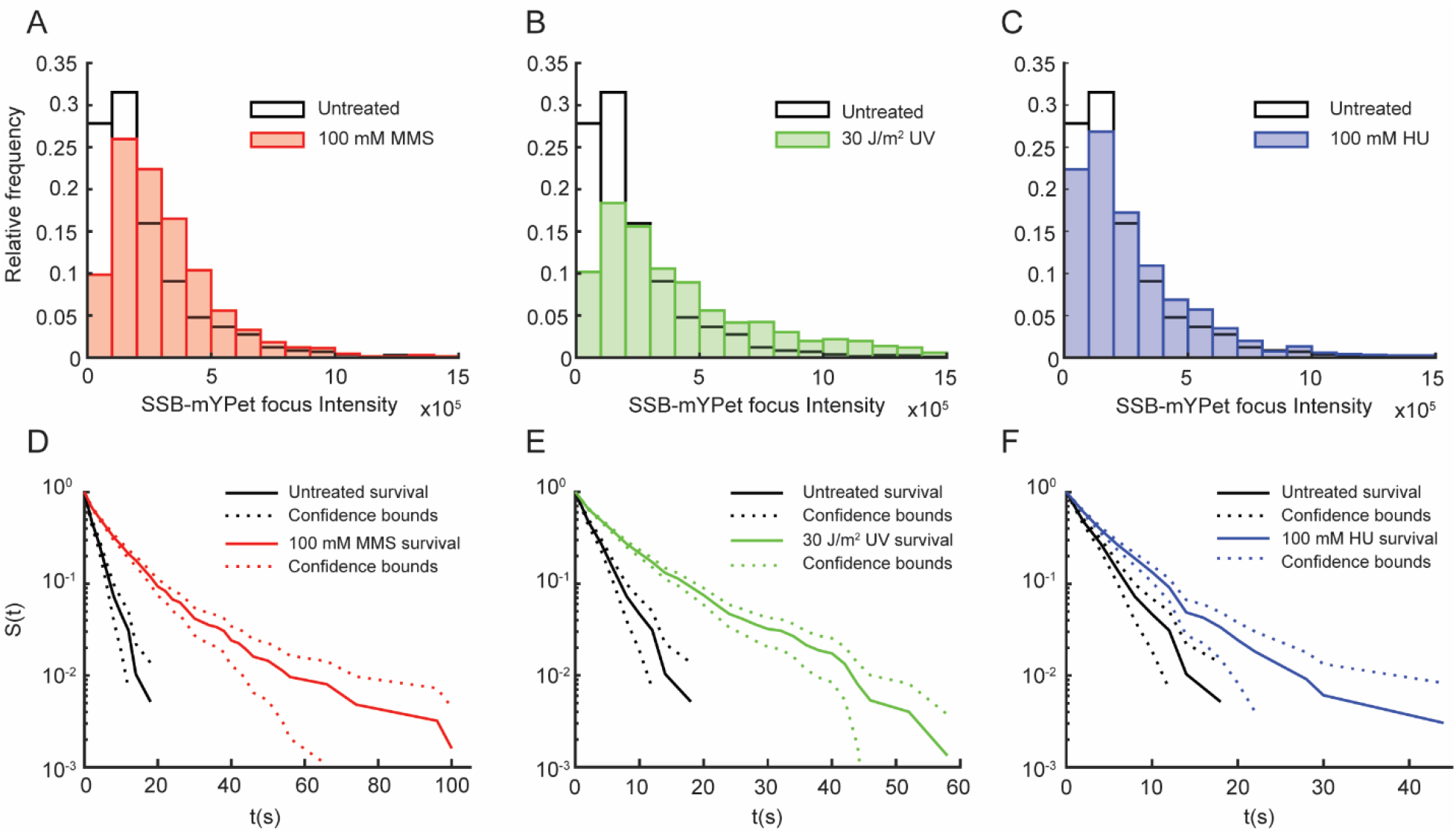
Effect of MMS, UV, and HU treatment on SSB-mYPet focus intensity and survival time of static SSB-PAmCherry tracks. Distributions of integrated SSB-mYPet focus intensity in untreated cells (white, *N* = 2,138) and (*A*) cells treated with 100 mM MMS (red, *N* = 1,309), (*B*) 30 J/m^2^ UV light (green, *N* = 2,015), and (*C*) 100 mM HU (blue, *N* = 1,207). Raw survival curves of static SSB-PAmCherry tracks in untreated (black, *N* = 545) cells and (*D*) MMS-treated (red, *N* = 1,171), (*E*) UV-treated (green, *N* = 1,755), and (*F*) HU-treated (blue, *N* = 956) cells measured with a 2-s stroboscopic interval.

To determine if the SSB binding lifetime also increased upon treatment, we imaged single SSB-PAmCherry molecules using a long 250 ms integration time with a stroboscopic illumination pattern of one excitation frame every two seconds; this stroboscopic illumination allowed us to extend the time until PAmCherry photobleaching and thus probe longer binding events. The resulting distributions of SSB binding lifetimes for untreated cells and cells treated with 100 mM MMS, 30 J/m^2^ UV, and 100 mM HU (Figure S4 and Table S6) were converted to survival curves (see Figure S5, Table S7, and Methods). The resulting survival curves (Figure 4D – F) revealed an increase in the SSB binding lifetime upon perturbations to replication, apparent as an increase in the population of molecules with longer binding times and a concomitant decrease in the population with shorter binding times. To quantify these results, each survival curve was fit to a double exponential function and corrected for PAmCherry photobleaching (see Figures S5 and S6, Table S8, and Methods). The lifetime increased from 4.5 s in untreated cells to 32.5 s, 56.7 s, and 8.9 s in cells treated with MMS, UV, and HU respectively, indicating a substantial increase in the SSB binding lifetime upon replication perturbation. Directly measured lifetime distributions give qualitatively similar results (Figures S4 and S5 and Table S6).

In addition to this increase in the binding lifetime of SSB molecules, we observed an increase in the number of static SSB molecules using these imaging conditions (Figure S7 and Table S9). The average number of SSB binding events increased by approximately two-fold for MMS, three-fold for UV treatment, and about 20% for HU treatment in the 2-second stroboscopic condition (Figure S7 and Table S9). Although these measurements do not report directly on the copy number of SSB at the replication fork, because they do not include molecules that are bound only briefly, the behavior is consistent with an increase in the static, or more stable, population of SSB.

### Other SSB interacting proteins are enriched at replication forks in the presence of DNA damage

Taken together, our results show that Pol IV is enriched at replication forks when replication is perturbed, primarily through interactions with SSB, and that this enrichment is likely a consequence of changes in SSB copy number and binding lifetime upon replication stalling. Importantly, this enrichment occurs even when Pol IV is unable to resolve the stall by performing TLS. In addition to Pol IV, the C-terminal tail of *E. coli* SSB is known to interact with at least 17 other SIPs(16, 18) raising the possibility that selective enrichment to stalled replication forks is general rather than specific to Pol IV. To test this possibility, we created C-terminal fusions of PAmCherry to two other SIPs, PriA and RecG, at their native loci. PriA is a helicase and a member of the primosome that plays a role in replication initiation and in the restart of stalled replication forks.(39–42) RecG is a helicase that likewise plays a role in replication restart at stalled forks.(42–44)

The functionality of the PriA and RecG fusions was tested by assaying their sensitivity to MMS. The survival of the PriA-PAmCherry fusion strain was essentially identical to the untagged PriA strain, both in the MG1655 WT background and in the *lexA51* imaging strain background (Figure S1C). The RecG-PAmCherry fusion showed a modest reduction in survival at higher MMS concentrations compared to untagged RecG. Cells bearing a Δ*recG* knockout, however, were unable to tolerate even the lowest MMS concentration tested, indicating that the RecG-PamCherry fusion still retains substantial activity (Figure S1D).

Unperturbed *E. coli* cells bearing a PriA-PAmCherry or RecG-PAmCherry fusion and an SSB-mYPet replisome marker were imaged using 250 ms integration times to resolve static molecules. There was no enrichment of either PriA or RecG near sites of replication in the absence of damage (Figure 5A – B), as observed for Pol IV. Upon treatment with 100 mM MMS, however, both proteins were strongly enriched at replication forks, with an enrichment of approximately four-fold for PriA and ten-fold for RecG. These results suggest that the replication stalling-dependent enrichment of SIPs may be a general mechanism triggered by changes in SSB binding dynamics, which acts to recruit DNA repair and DNA damage tolerance proteins when replication is challenged.

**Fig. 5.**
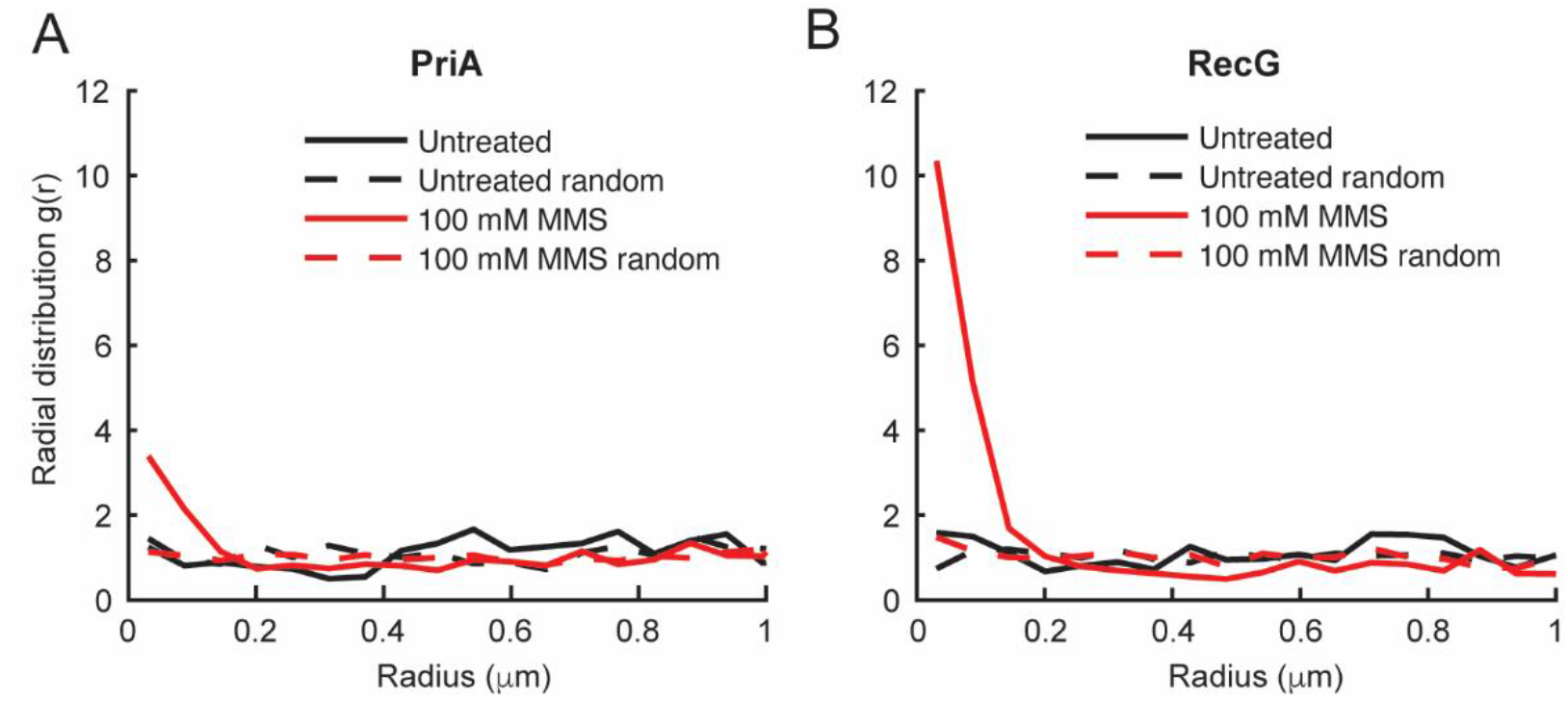
Effect of MMS treatment on the single-cell colocalization of SSB-interacting proteins PriA and RecG with SSB. (*A*) Radial distribution function *g*(*r*) between static PriA-PAmCherry molecules and SSB-mYPet foci in untreated cells (black, *N* = 781) and cells treated with 100 mM MMS (red, *N* = 2,412). (*B*) Radial distribution function *g*(*r*) between static RecG-PAmCherry molecules and SSB-mYPet foci in untreated (black, *N* = 1,189) and MMS-treated (red, *N* = 1,694) cells.

## Discussion

DNA damage acts to block the progress of the replisome. Numerous polymerases, helicases, and other enzymes not believed to be constitutive members of the replisome are employed to resolve these blockades. Previously we showed that the TLS polymerase Pol IV is enriched near replication forks upon treatment with the cognate DNA damaging agent MMS through interactions with the β clamp and SSB.(4, 17) Although MMS-induced lesions are a strong block for the replicative polymerase Pol III, Pol IV is able to bypass them efficiently.(31, 17) In this study we found that Pol IV is likewise selectively enriched when replication is perturbed by a non-cognate form of DNA damage, UV lesions, or by nucleotide depletion caused by HU or guanazole treatment. Thus, stable enrichment of Pol IV is not contingent on its ability to resolve the stall. Further, this enrichment requires the same β clamp and SSB interactions as in the case of MMS, with the SSB interaction playing the major role. These results are consistent with a model where Pol IV is rapidly enriched near the fork in response to general replication perturbations through the same interactions that mediate its response to cognate lesions.

Although interactions between Pol IV and the β clamp are essential for TLS, the interaction with SSB is critical for Pol IV to gain access to the clamp.(17) While clamps may reside on DNA behind the replication fork,(45) there is presumably only one clamp near a lesion site. Thus, binding sites on the clamp near a DNA lesion are limited. The β clamp is a dimer, with an equivalent binding cleft site on each protomer.(5) The Pol III α subunit binds tightly to one of these clefts, while the Pol III ε subunit binds weakly to the other cleft.(11, 46) Although this interaction has a lower binding affinity, ε is present at high effective local concentration due to α binding and acts as a molecular gate to regulate other factors from binding the clamp.(12) In addition, there are a number of other clamp-binding proteins that may compete with Pol IV for a cleft site. Thus, SSB acts as a platform that locally concentrates Pol IV near the clamp, allowing it to outcompete other clamp-binding proteins that do not bind SSB.(17)

SSB is a constitutive component of replication forks, where it binds ssDNA on the lagging strand. A previous imaging study revealed an average of approximately 8 SSB tetramers present at a single replication fork, a value consistent with typical Okazaki fragment sizes (650 – 2,000 bp depending on temperature)(47, 48) and SSB binding footprint (35 or 65 bp).(49, 16) Thus, some mechanism must prevent Pol IV enrichment through the interaction with SSB during processive replication. In this study, we looked for changes in SSB behavior upon replication perturbation that might explain this selective Pol IV enrichment. By directly imaging single SSB molecules and characterizing their diffusion, we identified both static (DNA-bound) and mobile populations. As expected, static SSB molecules were preferentially enriched near replication forks. Upon different forms of replication perturbation, we observed a significant shift in SSB dynamics, with the static fraction roughly doubling from 25% to 50% of the total SSB population. We propose that this change in SSB dynamics could represent a damage-dependent signal for Pol IV enrichment, explaining why Pol IV is not enriched near replication forks in the absence of replication stress. Although the SSB diffusion measurements are indicative of an increase in the total number of statically-bound SSB molecules, they are convoluted with a second possible effect, a change in SSB binding lifetime upon replication perturbation. We explored both of these possible mechanisms in this study.

The effect of replication perturbations on the amount of exposed ssDNA, and thus on the copy number of bound SSB, likely depends on the nature of the perturbation, and the effects may be different on the leading and lagging strand. During normal replication, there is no significant SSB binding on the leading strand due to the continuous nature of leading-strand synthesis (Figure 6, left). Stalling of the leading-strand replicative polymerase at a lesion, however, leads to uncoupling of the polymerase and helicase, exposing ssDNA on the leading strand (Figure 6, bottom right).(50) Alternatively, the presence of lesions on either the leading or lagging strand may result in repriming, in which case an ssDNA gap remains behind the moving fork.(12, 51, 52) We would expect UV irradiation, which produces highly blocking DNA lesions, to lead to repriming and ssDNA gaps behind the fork.(51, 53, 50) Nucleotide depletion by HU could lead to polymerase-helicase uncoupling and the accumulation of ssDNA on the leading strand. (54, 55) By measuring the intensity of SSB-mYPet foci, we found a two-fold increase upon UV treatment, but smaller increases of 50% and 25% for MMS and HU treatment, respectively. The relatively modest increase in focus intensity upon MMS treatment in comparison to UV may reflect the increased propensity for Pol IV-mediated TLS at the replication fork versus resolution by repriming. Despite the relatively modest increase in bound SSB for HU treatment, there was still a significant enrichment of Pol IV under this condition, suggesting that SSB copy number alone is not likely to be the sole determinant of Pol IV enrichment and that other factors, like the stability of Pol IV at the primer-template junction, may contribute. Because SSB clustering is needed to overcome the weak affinity of a single Pol IV-SSB C-terminal tail interaction,(17) an increase in the copy number of bound SSB may be required for Pol IV enrichment upon replication stalling.

**Fig. 6.**
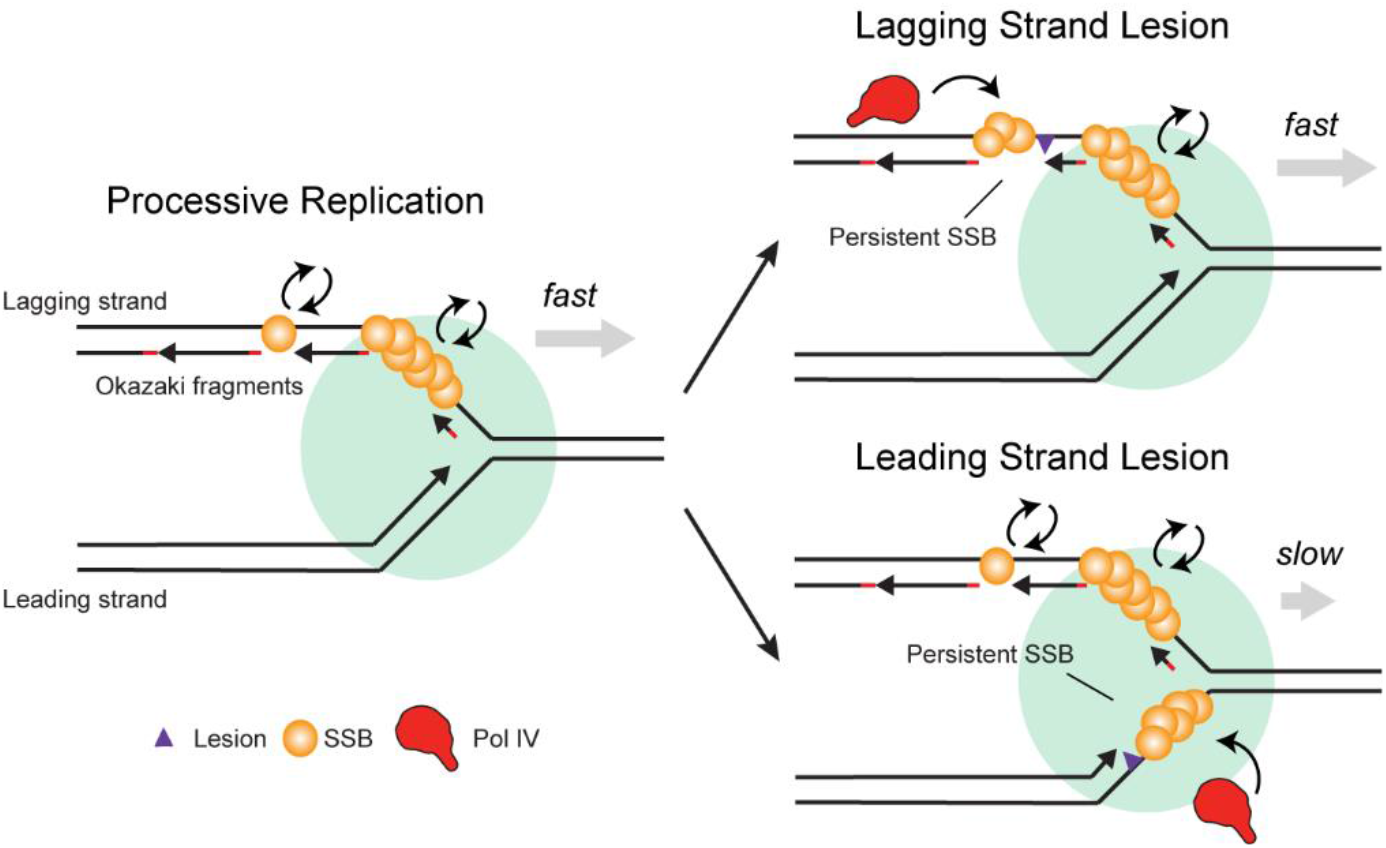
Model of the selective recruitment of Pol IV and other SIPs to stalled replication forks by SSB. During processive replication (left), SSB turns over rapidly on the lagging strand. In the case of a lagging strand lesion (top right), SSB bound at persistent lagging-strand ssDNA gaps enriches Pol IV or other SIPs. In the case of a leading strand lesion (bottom right), continued helicase translocation after Pol III stalling creates a persistent leading-strand ssDNA; SSB binding at this gap enriches Pol IV or other SIPs.

Like SSB copy number, the average binding lifetime of SSB may also change upon replication perturbation. During processive replication, the lifetime of an SSB molecule on the lagging strand is likely limited by eviction due to the synthesizing polymerase (Figure 6, left). Indeed, we measured an SSB lifetime of approximately 4 seconds (Table S8), consistent with estimated replication speeds of 500 – 1,000 base pairs per second and Okazaki fragments of up to 2,000 base pairs in length. Recycling of SSB from one Okazaki fragment to the next would increase the measured binding lifetime.(56) However, this is probably not the predominant mechanism in cells given that high in vivo SSB concentrations (300 – 600 nM) favor free SSB binding from solution.(56) Stalling at lesions on the lagging strand template is believed to be resolved by rapid repriming, resulting in persistent gaps and SSB stabilization (Figure 6, top right).(57) Likewise, SSB bound in gaps downstream of a stalled leading-strand polymerase should remain bound longer than lagging-strand SSB during processive replication (Figure 6, bottom right). Consistent with our expectations, perturbing replication resulted in increases in SSB binding lifetime from two-fold to greater than ten-fold. Thus, this reduction in turnover of bound SSB could lead to a corresponding reduction in Pol IV turnover and, as a result, an increase in average Pol IV enrichment.

We provide evidence that changes in SSB copy number and dynamics alone may trigger selective recruitment of Pol IV upon replication fork stalling. Other mechanisms are possible, however, either independent of the changes we observe or downstream of them. For example, post-translational modifications of SSB triggered by perturbations to replication could increase its affinity for Pol IV; in eukaryotes, post-translational modifications to the ssDNA-binding protein Replication Protein A (RPA) have been shown to play a role in the DNA damage response.(58) However, at present there is no evidence for such modifications in *E. coli*. Alternatively, a change in SSB binding conformation in response to stalling could result in selective enrichment. DNA binding is known to release the C-terminal tail of SSB, which can interact with the OB fold domain in free SSB, making it available for interaction with SIPs.(59) Further conformational changes in SSB upon replication fork stalling are not known, however, and proposals that SIP binding remodels the SSB filament(60, 61) do not explain how SSB selectively enriches factors at stalled forks. Recent observations show that SSB readily forms phase separated condensates in vitro.(62, 63) These condensates may preferentially form at clusters of persistent SSB, thus leading to selective enrichment of SIPs at stalled replication forks. However, further work is needed to determine if SSB condensates are physiologically relevant and whether their formation is required for SIP recruitment in cells.

In *E. coli* there are at least 17 other SIPs that are involved in various genome maintenance processes. Among these are RecG and PriA, which help resolve stalled replication forks. Prior studies have not found strong evidence for the formation of replisome-associated foci of RecG(64–66) or PriA(64, 67, 65, 66) at native expression levels during processive replication, nor have they investigated whether these SIPs are selectively enriched near replication forks upon replication perturbation. To address these questions, we imaged PriA and RecG with single-molecule resolution and quantified their colocalization with sites of replication. We find that both PriA and RecG are strongly enriched near replication forks upon MMS treatment, but not in unperturbed cells, similar to Pol IV. Although the SSB binding interfaces are not known for these proteins, and thus we cannot make mutations to eliminate binding, it is likely that this enrichment is mediated through interactions with SSB. Previously, we showed that a chimeric protein composed of the C-terminal Pol IV little finger domain and the RecQ SSB-binding winged helix domain was selectively enriched upon DNA damage, and that this enrichment was lost when the SSB-binding residues of RecQ were mutated.(17) As for Pol IV, this selective enrichment in response to replication stress likely helps PriA and RecG gain access to the DNA when their activity may be needed. Our results suggest that selective enrichment of SIPs in response to replication perturbations is a general effect, but future work is needed to elucidate whether other interactions help to establish a hierarchical response of different SIPs upon replication stalling and to determine the functional consequences for cell survival upon replication perturbation.

## Methods

### Bacterial strain construction

Bacterial strains containing fluorescent protein fusions were constructed using Lambda Red recombineering(68) and P1*vir* transduction as previously described.(4) In brief, the *E. coli* K-12 type strain, MG1655, was transformed with the temperature-sensitive recombineering plasmids pSIM5 (chloramphenicol) or pSIM6 (ampicillin).(69) Recombineering fragments bearing approximately 50 bp length homology arms were generated by PCR amplification (Q5 polymerase, New England Biolabs) of pKD3- or pKD4-based plasmids and transformed into MG1655 pSIM5 or pSIM6 by electroporation. Recombinants were selected on LB agar plates containing the appropriate antibiotic (chloramphenicol for pKD3, kanamycin for pKD4) and the recombineering plasmids were cured by growth at 37 °C. The modified alleles were then moved into the appropriate strain background for imaging using P1*vir* transduction. When required for further strain construction steps, FRT-flanked antibiotic cassettes were removed using Flp-FRT recombination with the temperature-sensitive plasmid pCP20, leaving a scar of approximately 80 bp containing a single FRT site.(70) Fusions were validated at each step by PCR amplification of genomic DNA and Sanger DNA sequencing. All oligonucleotides, plasmids, and bacterial strains used in this study are listed in Tables S10 – S12 and construction details for individual strains are listed in the Supplementary Methods.

The photoactivatable fluorescent protein PAmCherry1(19) was used to make Pol IV, SSB, PriA, and RecG fusions for PALM imaging, and the monomeric YFP variant mYPet was used to make SSB and e replisome marker fusions. All fusions contained a linker between the C-terminus of the protein and the N-terminus of the fluorescent protein: (GGGS)4 for PamCherry fusions to Pol IV, PriA, and RecG; SSAGSAAGSGEF for the mYPet and PAmCherry fusions to SSB; and SAGSAAGSGEF for the mYPet fusion to ε.

### Culture conditions and sample preparation for microscopy

Cell culture and sample preparation methods were as previously described.(4) In brief, cells were streaked from glycerol stocks onto LB agar plates containing the appropriate antibiotics and incubated overnight at 37 °C. Single colonies were picked and used to inoculate “overday” starter cultures in 3 mL LB, which were incubated for approximately 8 h on a roller drum at 37 °C. Overnight cultures in 3 mL supplemented M9 glucose medium (0.4% glucose, 1 mM thiamine hydrochloride, 0.2% casamino acids, 2 mM MgSO_4_, and 0.1 mM CaCl_2_) were inoculated with a 1:1,000 dilution of the overday cultures and then grown in a roller drum at 37 °C overnight. The next day, imaging cultures were prepared in 50 mL volume of supplemented M9 medium, inoculated with a 1:200 dilution of the corresponding overnight cultures, and incubated at 37 °C shaking at 225 rpm. Antibiotics were not included in liquid cultures, but 0.5 mM IPTG was included in the M9 overnight and imaging cultures to induce expression of fusions inserted at the *lacZ* locus.

When imaging cultures reached early exponential phase (OD_600nm_ ≈ 0.15), a 1 mL aliquot was removed and centrifuged at 8,609 × *g*. The supernatant was removed, the cell pellet was resuspended in a few μL of the remaining liquid, and < 1 μL was deposited on an agarose pad (3% concentration of NuSieve GTG agarose) prepared with supplemented M9 glucose medium (not including thiamine hydrochloride, casamino acids, or IPTG). The agarose pad was sandwiched between two glass coverslips. Coverslips were cleaned by 30 min cycles of sonication in ethanol and then 1 M KOH (two cycles of each, alternating) and rinsed thoroughly with DI water.

### Sensitivity assays and treatment conditions for microscopy

A spot dilution assay was used to determine cell sensitivity to MMS and HU. In brief, 50 mL LB cultures were inoculated with a 1:1,000 dilution of the corresponding overnight cultures and incubated shaking at 37 °C until reaching OD600nm ≈ 1.0. Aliquots were removed and serially diluted in 0.9% NaCl. A dilution series was stamped or pipetted on LB agar plates containing MMS (2, 4, or 6 mM), HU (4, 7, and 10 mM), or without any drug. All plates included 0.5 mM IPTG to induce SSB expression. Plates were photographed after a 16 h incubation at 37 °C.

As described previously,(4, 26) cells were treated with methyl methanesulfonate (MMS) included at 100 mM concentration in the standard agarose pad. Cells were deposited on the pad as normal, then incubated at room temperature in a humidified chamber for 20 min before imaging. Cells were treated with 100 mM hydroxyurea (HU) for 20 min following the same approach. UV treatment was performed using 254 nm light (General Electric G15T8 15W bulb) at an irradiance of 2 W/m^2^ measured with a UVP UVX Radiometer (#97-0015-02) and UVX-25 sensor. Cells were deposited on an agarose pad as normal, but without the coverslip in place, and then exposed to a fluence of 30 J/m^2^. After treatment, the coverslip was placed on the agarose pad and the sample was imaged immediately.

### Microscopy

The fluorescence microscope used in this study was described previously.(4) In brief, a Nikon TE2000 inverted microscope was equipped with a 514 nm laser (Coherent Sapphire, 150 mW) for mYPet excitation, a 405 nm laser (Coherent OBIS, 100 mW) for PAmCherry activation, and a 561 nm laser (Coherent Sapphire, 200 mW) for PAmCherry excitation. The lasers were passed through excitation filters (Chroma ZET405/20X, ZET514/10X, and ZET561/10X), and combined with dichroic filters (Chroma ZT405rdc, Chroma ZT514rdc, and a mirror). Imaging was performed using highly inclined thin illumination,(71) or near-TIRF, in which a 400 mm focal length lens was used to focus the beams to the back focal plane of a Nikon CFI Apo 100×/1.49 NA TIRF objective. Two-color imaging was performed using a Chroma 91032 Laser TIRF Cube containing a ZT405/514/561rpc dichroic filter, ZET442/514/561m emission filter, and ET525lp longpass filter. Images were collected on a Hamamatsu ImageEM C9100-13 EMCCD camera. A brightfield image was recorded for each field of view using white light transillumination.

Movies were recorded with an integration time of 13.3 ms for short exposure imaging and 250 ms for long exposure imaging, as described previously.(4) Stroboscopic imaging was implemented using computer-controlled shutters (Uniblitz VS14) and custom-written LabVIEW (National Instruments) scripts. All short and long exposure PALM movies initiated with a 561 nm pre-bleaching period, followed by 514 nm excitation to image mYPet fusions, followed by simultaneous low-power 405 nm excitation and 561 nm excitation for photoactivation and imaging of PAmCherry fusions. The 405 nm power was increased twice during the course of the PALM movie. Laser power densities at the sample were as follows: ~ 120 W/cm^2^ (short exposure) or ~ 12.5 W/cm^2^ (long exposure) for 561 nm excitation; ~ 0.23 W/cm^2^ (SSB-mYPet for long exposure focus intensity analysis to avoid focus saturation), ~ 0.4 W/cm^2^ (SSB-mYPet in other long exposure imaging), 1.4 W/cm^2^ (ε-mYPet long exposure), or 16.6 W/cm^2^ (ε-mYPet short exposure) for 514 nm excitation; and ~ 2.5 – 17.5 mW/cm^2^ 405 nm excitation.

### Image analysis

Automated image analysis was carried out in MATLAB using the approach and parameters described in detail previously.(4) In brief, the MicrobeTracker(72) and u-track(73, 74) packages were used for cell segmentation of brightfield images and fluorescence spot detection and tracking, respectively. Fluorescent spots were fit to symmetrical 2D Gaussian point spread functions (PSFs). Static molecules in long exposure PALM imaging were identified by comparing the average PSF width over all localizations in the track to the mean PSF width measured in fixed cells.(4) SSB-mYPet and ε-mYPet foci were fit to 2D Gaussian PSFs after averaging the first 5 frames of 514 nm excitation; to avoid spurious detection of broad and weak fluorescence spots, foci with background values below the camera offset level were discarded.

### Data analysis

Data analysis methods were as described in detail previously.(4)

#### Diffusion coefficient analysis

Apparent two-dimensional diffusion coefficients (*D*\*) were calculated for short exposure PALM imaging. First the mean-squared displacement was determined for tracks with a number of localizations *N* ≥ 5 as:

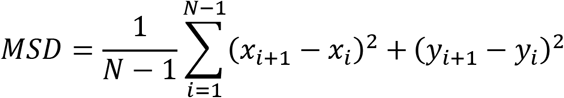

where *x* and *y* are the track coordinates. Then *D*\* was calculated as:

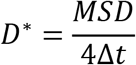

where Δ*t* is the time interval between localizations in the track.

#### Diffusion coefficient distribution fitting

Probability distributions of apparent distribution coefficients were fit to an analytical expression for three diffusing species where trajectories contain exactly four steps(75):

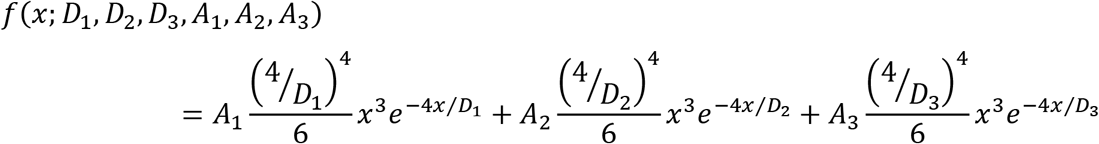

with the constraint:

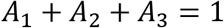

Here *A*_n_ represents the fraction of molecules corresponding to the *n*^th^ species and *D*_n_ represents the diffusion coefficient of the *n*^th^ species. For this analysis, trajectories longer than four steps were truncated and the apparent diffusion calculated as above.

#### Radial distribution function analysis

Colocalization was quantified using radial distribution function analysis,(29, 30) as described previously.(4) In brief, a distribution was generated of the mean distance between each Pol IV-PAmCherry trajectory within a cell and the nearest SSB-mYPet focus. Then a random Pol IV-SSB distance distribution was generated by taking the same cell outline and SSB focus position(s) and simulating the same number of Pol IV localizations randomly across the cell. The same procedure was repeated for all cells in the dataset to yield aggregated experimental and simulated Pol IV-SSB distance distributions. Finally, the radial distribution function *g*(*r*) was calculated by normalizing the experimental distance distribution by the simulated one. As described previously, 100 different simulated random distributions were generated to account for variability, giving 100 *g*(*r*) curves. The final *g*(*r*) curve was taken as the mean of these 100 curves. Independently, another random distance distribution was simulated and normalized by the same 100 simulated random distributions to give a mean random *g*(*r*) curve; deviations in this random *g*(*r*) curve from 1 may arise due to the finite sample size. This procedure was repeated for colocalization analysis of other PAmCherry-labeled proteins and mYPet-labeled replisome markers.

Survival curve analysis: The empirical cumulative distribution function, *F*(*t*), with 95% confidence bounds was tabulated for each set (untreated, MMS-damaged, UV-damaged, and HU-treated) of SSB-PAmCherry molecule lifetimes using the MATLAB ecdf function. The survivor function, the complement, was then tabulated using the relationship *S*(*t*) = 1 – *F*(*t*). For these analyses, only events lasting at least two frames were considered to reduce bias introduced by nonspecific binding that may be captured in a single frame. The result was fit to a normalized double exponential:

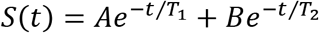

with the requirement that *A* + *B* = 1. The quantity *A*×*T_1_* + *B*×*T_2_* represents the weighted estimated survival time *T* of SSB-PAmCherry.

#### Photobleaching analysis

The survival analysis described above was generated for continuous imaging as well as 1-s and 2-s stroboscopic imaging intervals. As described previously,(76, 77) this allows us to plot the measured off-rate of any given SSB-PAmCherry molecule as a function of total stroboscopic interval. Here, we assume that while the off-rate of the molecule from the replisome is a constant, the off-rate contribution from photobleaching is weighted by the total laser exposure within that time interval, following the expression below.

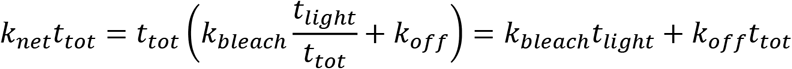

In the expression above, *k*_net_ is the inverse of *T*, or the measured weighted survival time of the SSB-PAmCherry molecule. For this analysis, we included all events and fit the data with a double-exponential function (Figure S5). We used a net survival time *T* as a sum of the two survival times weighted by their pre-exponential factors as the treatment condition-dependent value. The net observed off-rate is the sum of the rate of photobleaching and the off-rate of the particle itself, but the photobleaching rate is weighted by *t*_light_/*t*_tot_, where *t*_light_ is the exposure time of the light frame and *t*_tot_ is the duration of the total stroboscopic interval (including light and dark frames). Thus, when plotting *k*_net_*t*_tot_ vs. *t*_tot_, the intercept is *k*_bleach_*t*_light_, or the expected number of photobleaching events per light frame, and the slope is *k*_off_, an estimated value for the off-rate of the fluorescent molecule corrected for the impact of photobleaching (Figure S6).

#### Statistical analysis

The two-sided Wilcoxon rank-sum test (MATLAB function ranksum) was used to compare different distributions, with statistically significant differences determined as *p* < 0.05.

## Data and code availability

The raw data and analysis code from the current study are available from the corresponding authors upon request.

## Acknowledgments

We thank members of the Loparo lab for helpful feedback. We also thank Talley Lambert and the Nikon Imaging Center at Harvard Medical School for help with the collection of preliminary data. This work was supported by National Institutes of Health grants R01 GM114065 (to J.J.L.), F32 GM113516 (to E.S.T.), and T32 GM008313 (to S.C.P.), as well as a Landry Cancer Biology Research Fellowship (to S.C.P.).

