## Supplementary Information for "“Replication stalling activates SSB for recruitment of DNA damage tolerance factors”"

**This PDF file includes:**

Figures S1 to S7
Tables S1 to S12
Supplementary Methods
Supplementary References

### Supplementary Figures

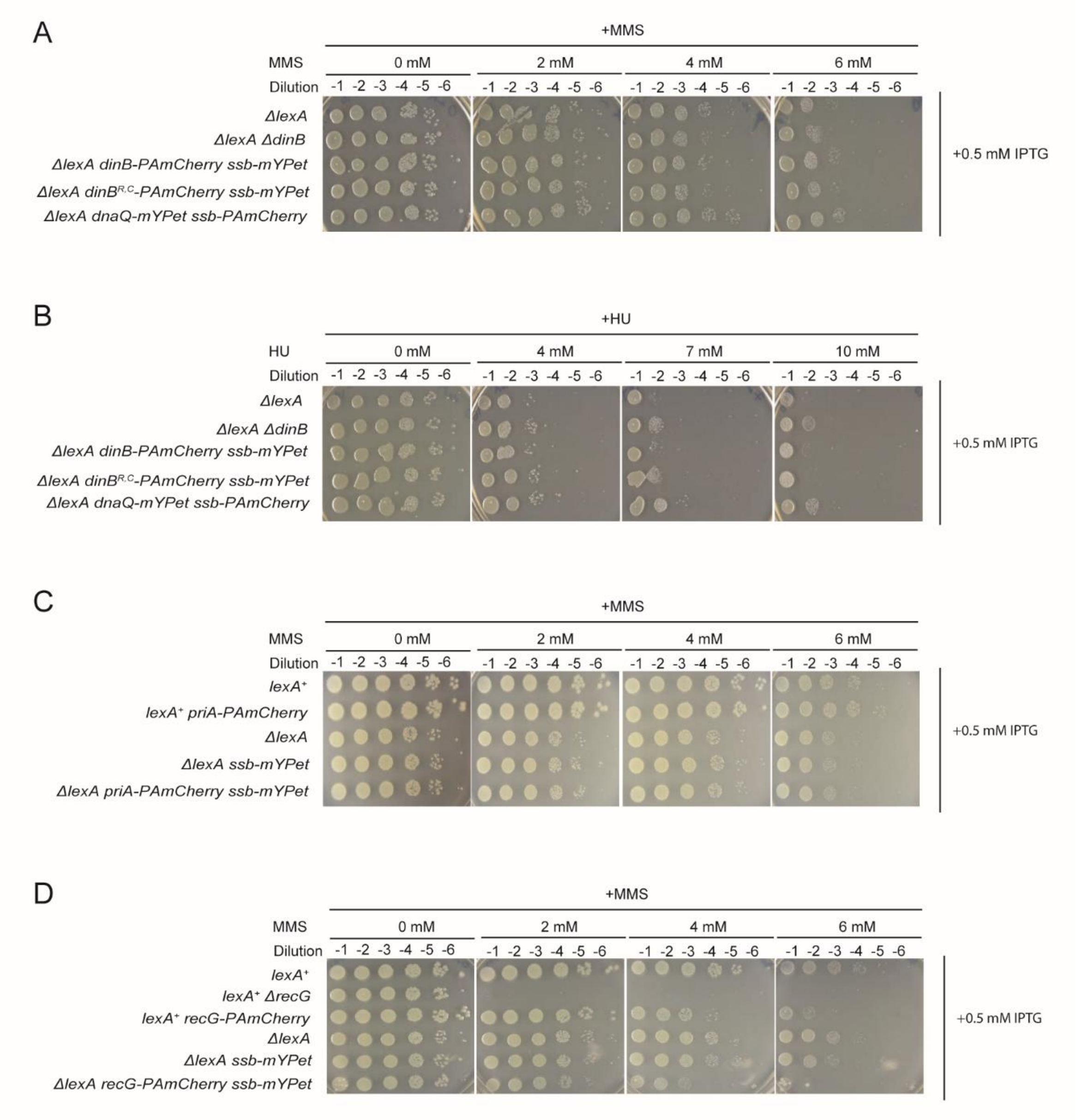


**Fig. S1.** (*A*) Serial 10-fold dilutions of *E. coli* strains with mutants and fusions to Pol IV (*dinB*) grown on LB agar plates with and without methyl methanesulfonate (MMS) added at 0, 2, 4, and 6 µM concentration. (*B*) Serial 10-fold dilutions of *E. coli* strains with mutants and fusions to Pol IV (*dinB*) grown on LB agar plates with and without hydroxyurea (HU) added at 0, 4, 7, and 10 µM concentration. Serial 10-fold dilutions of *E. coli* strains with fusions to (*C*) PriA (*priA*) or (*D*) RecG (*recG*) grown on LB agar plates with and without MMS added at 0, 2, 4, and 6 mM concentration.

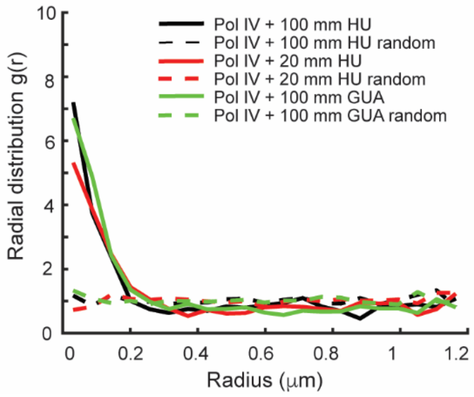


**Fig. S2.** Single-cell colocalization of Pol IV-PAmCherry and SSB-mYPet in ribonucleotide reductase inhibitor-treated cells. Radial distribution function *g(r)* between each static Pol IV-PAmCherry track and the nearest SSB-mYPet focus for cells treated for 20 min with 100 mM HU (black, *N* = 1,539), 20 mM HU (red *N* = 2,267), and 100 mM guanazole (green, *N* = 878).

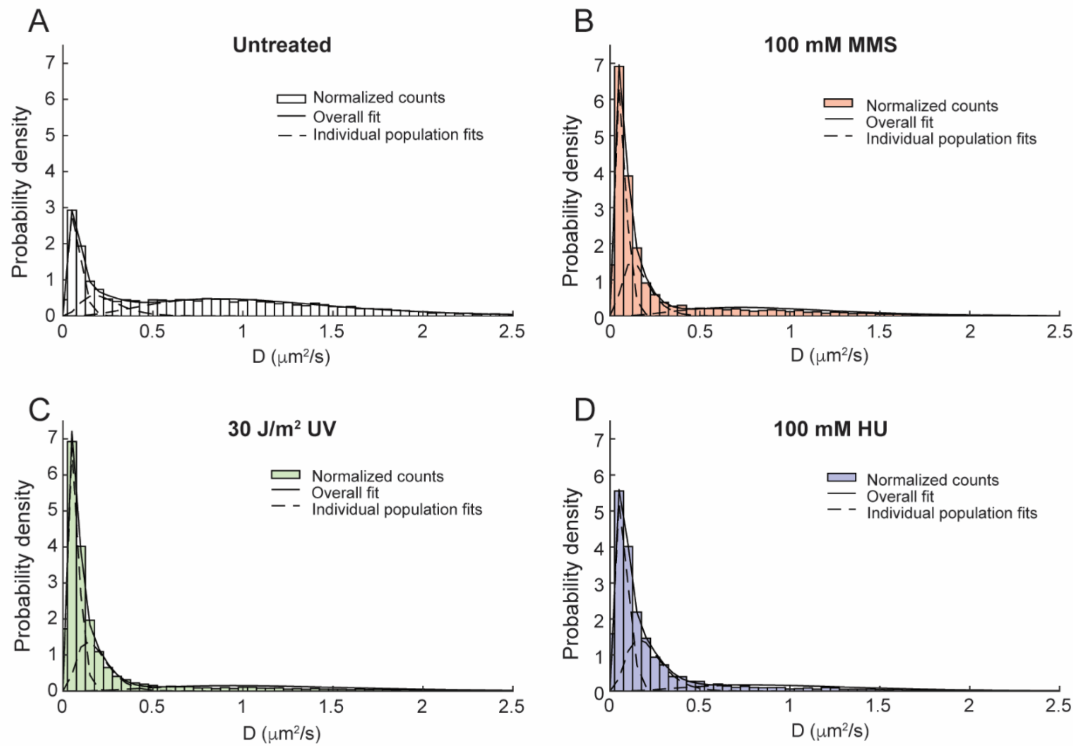


**Fig. S3.** Population fits to the distributions of the apparent diffusion coefficient (*D**) for SSB-PAmCherry in (*A*) untreated cells (*N* = 9,251) and cells treated with (*B*) 100 mM MMS (red, *N* = 11,797), (*C*) 30 J/m$^2$ UV light (green, *N* = 16,542), and (*D*) 100 mM HU (blue, *N* = 5,690). See Methods and Table S6 for fit equations and parameters.

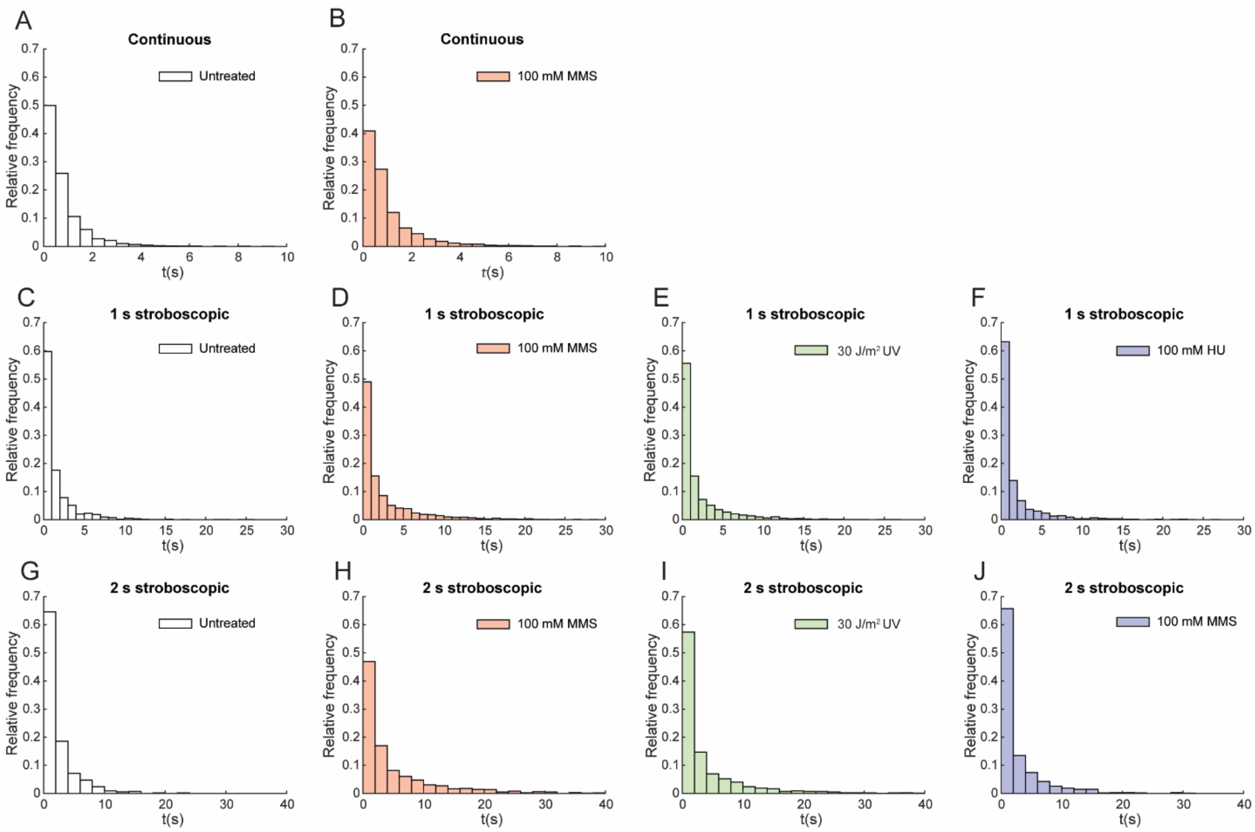


**Fig. S4.** Lifetime distributions of static SSB-PAmCherry tracks. Continuous imaging in (*A*) untreated cells (white, *N* = 2,965) and (*B*) cells treated with 100 mM MMS (red, *N* = 6,924) cells. Stroboscopic imaging with a 1-s interval in (*C*) untreated cells (white, *N* = 1,403) and cells treated with (*D*) 100 mM MMS (red, *N* = 3,837), (*E*) 30 J/m$^2$ UV light (green, *N* = 2,922), and (*F*) 100 mM HU (blue, *N* = 1,659) cells. Stroboscopic imaging with a 2-s interval in (*G*) untreated cells (white, *N* = 545) and cells treated with (*H*) 100 mM MMS (red, *N* = 1,171), (*I*) 30 J/m$^2$ UV light (green, *N* = 1,755), and (*J*) 100 mM HU (blue, *N* = 956).

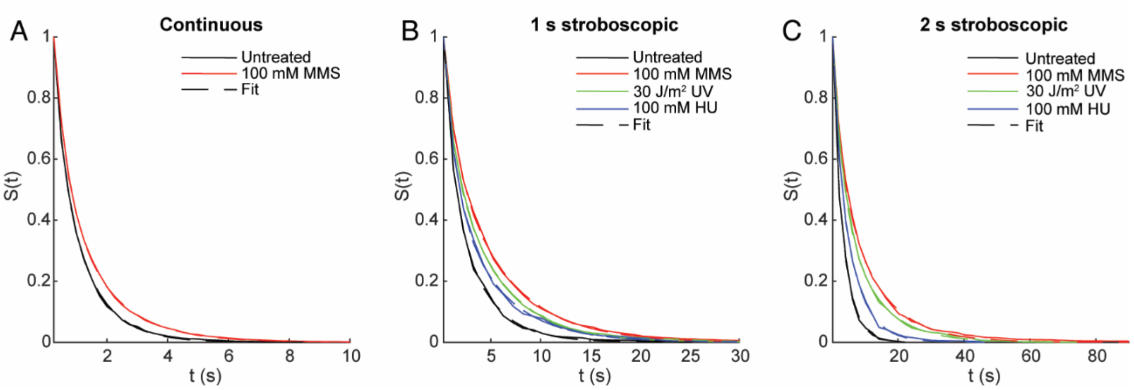


**Fig. S5.** Survival curves of static SSB-PAmCherry tracks fit to double-exponential decay functions. (*A*) Continuous imaging in untreated cells (white, $N = 2{,}965$) and cells treated with 100 mM MMS for 20 min (red, $N = 6{,}924$). (*B*) Stroboscopic imaging with a 1-s interval in untreated (white, $N = 1{,}403$) cells and cells treated with 100 mM MMS (red, $N = 3{,}837$), 30 J/m$^2$ UV light (green, $N = 2{,}922$), and 100 mM HU (blue, $N = 1{,}659$). (*C*) Stroboscopic imaging with a 2-s interval in untreated cells (white, $N = 545$) and cells treated with 100 mM MMS (red, $N = 1{,}171$), 30 J/m$^2$ UV light (green, $N = 1{,}755$), and 100 mM HU (blue, $N = 956$).

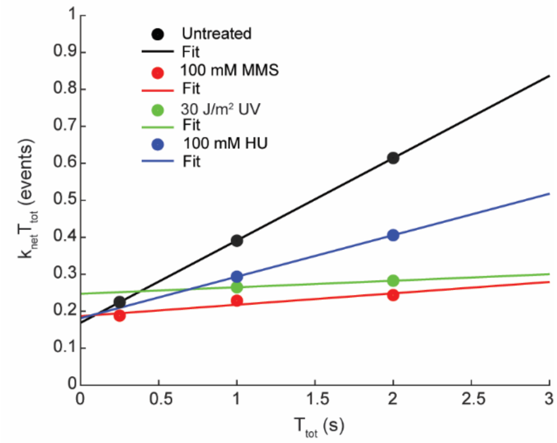


**Fig. S6.** Photobleaching correction to estimated SSB-PAmCherry binding lifetimes. Measured apparent dissociation rate constants $k_{net}$ from exponential fits to survival curves (see Figure S5) multiplied by the stroboscopic interval $T_{tot}$ plotted against $T_{tot}$ for untreated cells (black circles) and cells treated with 100 mM MMS (red circles), 30 J/m$^2$ UV light (green circles), and 100 mM HU (blue circles). Linear fits (solid lines) for each condition give the photobleaching-corrected SSB-PAmCherry off rate as the slope and the PAmCherry photobleaching rate as the intercept (see Methods).

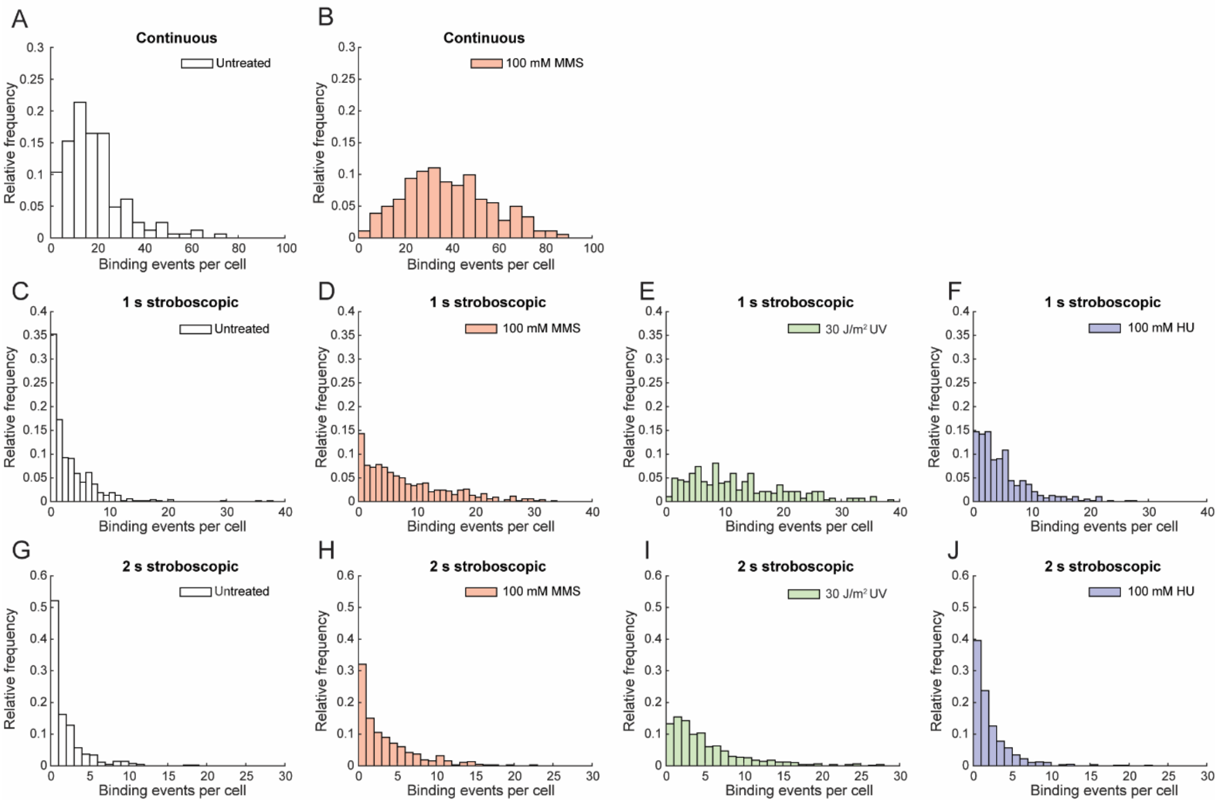


**Fig. S7.** Binding number distributions of static SSB-PAmCherry molecules for continuous imaging in (*A*) untreated cells (white, *N* = 164) and (*B*) cells treated with 100 mM MMS (red, *N* = 181). Stroboscopic imaging with a 1-s interval in (*C*) untreated cells (white, *N* = 539) and cells treated with (*D*) 100 mM MMS (red, *N* = 538), (*E*) 30 J/m$^2$ UV light (green, *N* = 285), and (*F*) 100 mM HU (blue, *N* = 387). Stroboscopic imaging with a 2-s interval in (*G*) untreated cells (white, *N* = 437) and cells treated with (*H*) 100 mM MMS (red, *N* = 380), (*I*) 30 J/m$^2$ UV light (green, *N* = 512), and (*J*) 100 mM HU (blue, *N* = 581).

### Supplementary Tables

#### Table S1: Estimated SSB-PAmCherry diffusion coefficients

| **Treatment condition** | **Mean ± S.E.M. (μm²/s)** | **Median (μm²/s)** | ***N* (events)** | ***N* per cell** | **Wilcoxon rank-sum *P*-value relative to untreated** |
|---|---|---|---|---|---|
| Untreated | 0.743 ± 0.007 | 0.638 | 9,251 | 5.1 | - |
| 100 mM MMS | 0.266 ± 0.004 | 0.095 | 11,797 | 12.9 | 0.00000 |
| 30 J/m² UV | 0.245 ± 0.003 | 0.094 | 16,542 | 23.2 | 0.00000 |
| 100 mM HU | 0.246 ± 0.005 | 0.118 | 5,690 | 6.8 | 0.00000 |

#### Table S2: Estimated SSB-PAmCherry diffusion coefficients sorted by distance from the replisome

| **Treatment condition** | ***N* (*D** < 0.25 μm²/s, static)** | ***N* (*D** > 0.25 μm²/s, mobile)** |
|---|---|---|
| Untreated (all) | 2,947 | 5,753 |
| Untreated (< 200 nm replisome) | 1,014 | 262 |
| Untreated (> 200 nm replisome) | 1,933 | 5,491 |
| 100 mM MMS (all) | 8,763 | 2,681 |
| 100 mM MMS (< 200 nm replisome) | 4,405 | 371 |
| 100 mM MMS (> 200 nm replisome) | 4,358 | 2,310 |
| 30 J/m² UV (all) | 12,268 | 3,077 |
| 30 J/m² UV (< 200 nm replisome) | 2,771 | 232 |
| 30 J/m² UV (> 200 nm replisome) | 9,497 | 2,845 |
| 100 mM HU (all) | 3,854 | 1,201 |
| 100 mM HU N (< 200 nm replisome) | 511 | 84 |
| 100 mM HU (> 200 nm replisome) | 3,343 | 1,117 |

#### Table S3: SSB-PAmCherry diffusion coefficient distribution fit parameters

| **Treatment condition** | **$D_1$ (μm²/s)** | **$A_1$** | **$D_2$ (μm²/s)** | **$A_2$** | **$D_3$ (μm²/s)** | **$A_3$** |
|---|---|---|---|---|---|---|
| Untreated | 0.0752 | 0.242 | 0.236 | 0.149 | 1.150 | 0.609 |
| 100 mM MMS | 0.0664 | 0.470 | 0.163 | 0.274 | 0.972 | 0.256 |
| 30 J/m² UV | 0.0689 | 0.501 | 0.184 | 0.287 | 1.256 | 0.212 |
| 100 mM HU | 0.0756 | 0.452 | 0.213 | 0.346 | 1.069 | 0.202 |

### Table S4: Integrated SSB-mYPet focus intensity

| Treatment condition | Mean ± S.E.M. (AUe-5) | Median (AUe-5) | *N* (foci) | Wilcoxon rank-sum *P*-value relative to untreated |
|---|---|---|---|---|
| Untreated | 2.40 ± 0.05 | 1.61 | 2,138 | - |
| 100 mM MMS | 3.16 ± 0.07 | 2.53 | 1,309 | 0.00000 |
| 30 J/m$^2$ UV | 6.6 ± 0.2 | 3.57 | 2,015 | 0.00000 |
| 100 mM HU | 2.87 ± 0.08 | 2.04 | 1,207 | 0.00000 |

### Table S5: Number of SSB-mYPet foci per cell

| Treatment condition | Mean ± S.E.M. | Median | *N* (cells) | Wilcoxon rank-sum *P*-value relative to untreated |
|---|---|---|---|---|
| Untreated | 1.06 ± 0.02 | 1 | 2,021 | - |
| 100 mM MMS | 1.11 ± 0.02 | 1 | 1,182 | 0.00311 |
| 30 J/m$^2$ UV | 1.22 ± 0.02 | 1 | 1,646 | 0.00000 |
| 100 mM HU | 1.29 ± 0.03 | 1 | 938 | 0.00000 |

### Table S6: SSB-PAmCherry binding lifetimes

| Treatment condition (Stroboscopic interval) | Mean ± S.E.M. (s) | *N* (events) | *N* (cells) | Wilcoxon rank-sum *P*-value relative to untreated |
|---|---|---|---|---|
| Untreated (2 s) | 1.7 ± 0.1 | 545 | 437 | - |
| 100 mM MMS (2 s) | 5.2 ± 0.3 | 1,171 | 380 | 0.00000 |
| 30 J/m$^2$ UV (2 s) | 3.6 ± 0.2 | 1,755 | 512 | 0.00001 |
| 100 mM HU (2 s) | 2.3 ± 0.1 | 956 | 581 | 0.69151 |
| Untreated (1 s) | 1.41 ± 0.06 | 1,403 | 539 | - |
| 100 mM MMS (1 s) | 2.66 ± 0.07 | 3,837 | 538 | 0.00000 |
| 30 J/m$^2$ UV (1 s) | 2.08 ± 0.07 | 2,922 | 285 | 0.00003 |
| 100 mM HU (1 s) | 1.62 ± 0.08 | 1,659 | 387 | 0.29163 |
| Untreated (continuous) | 0.72 ± 0.02 | 2,965 | 164 | - |
| 100 mM MMS (continuous) | 0.94 ± 0.01 | 6,924 | 181 | 0.00000 |

### Table S7: SSB-PAmCherry survival times

| Treatment condition | $T_1$ (s) (2 s interval) | $T_2$ (s) (2 s interval) | $T_1$ (s) (1 s interval) | $T_2$ (s) (1 s interval) | $T_1$ (s) (continuous) | $T_2$ (s) (continuous) |
|---|---|---|---|---|---|---|
| Untreated | 0.44 | 3.35 | 0.85 | 3.25 | 0.27 | 1.01 |
| 100 mM MMS | 2.70 | 11.31 | 1.14 | 5.48 | 0.44 | 1.49 |
| 30 J/m$^2$ UV | 2.96 | 10.38 | 0.93 | 4.74 | - | - |
| 100 mM HU | 1.75 | 5.67 | 1.10 | 4.80 | - | - |

### Table S8: Photobleaching-corrected SSB-PAmCherry survival times

| Treatment condition | $k_{off}$ (s$^{-1}$) | $T_{off}$ (s) | $k_{bleach}$ (s$^{-1}$) |
|---|---|---|---|
| Untreated | 0.2227 | 4.5 | 0.6752 |
| 100 mM MMS | 0.0308 | 32.5 | 0.7473 |
| 30 J/m$^2$ UV | 0.0176 | 56.7 | 0.9889 |
| 100 mM HU | 0.1122 | 8.9 | 0.7253 |
| **Mean $k_{bleach}$** | | | 0.7842 |

### Table S9: Number of SSB-PAmCherry binding events

| Treatment condition (Stroboscopic interval) | Mean ± S.E.M. (events) | Median (events) | *N* (cells) | Wilcoxon rank-sum *P*-value relative to untreated |
|---|---|---|---|---|
| Untreated (2 s) | 1.4 ± 0.1 | 0 | 437 | - |
| 100 mM MMS (2 s) | 3.1 ± 0.2 | 2 | 380 | 0.00000 |
| 30 J/m$^2$ UV (2 s) | 4.7 ± 0.2 | 3 | 512 | 0.00000 |
| 100 mM HU (2 s) | 1.8 ± 0.1 | 1 | 581 | 0.00086 |
| Untreated (1 s) | 3.0 ± 0.2 | 1 | 539 | - |
| 100 mM MMS (1 s) | 7.6 ± 0.3 | 5 | 538 | 0.00000 |
| 30 J/m$^2$ UV (1 s) | 11.8 ± 0.5 | 10 | 285 | 0.00000 |
| 100 mM HU (1 s) | 4.6 ± 0.2 | 3 | 387 | 0.00000 |
| Untreated (continuous) | 18.1 ± 1.0 | 15 | 164 | - |
| 100 mM MMS (continuous) | 38.3 ± 1.4 | 37 | 181 | 0.00000 |

### Table S10: Oligonucleotides and Gene Blocks used in this study

| Number | Designation | Sequence (5′-3′) |
|---|---|---|
| oET110 | recG-G4S-PAmC-KI-for | CTGATAGAACGCTGGATGCCGGAGACGGAACGTTACTCGAATG CGGGTGGTGGTGGTTCTGGTGG |
| oET111 | recG-PAmC-KI-rev | GGAAGGTAGGGTAACCTGAAATGGCGGTCTTCTCACTGCCGCC TTTCTTATGAATATCCTCCTTAGTTCC |
| oET114 | priA-G4S-PAmC-KI-for | TCCCGTAAGGTGAAATGGGTGCTGGATGTTGATCCGATTGAGG GTGGTGGTGGTGGTTCTGGTGG |
| oET115 | priA-PAmC-KI-rev | GTGATGAATATTGAATTTTTCGATCCGCCTCGCATCGTGAGCG GTTCTTATGAATATCCTCCTTAGTTCC |
| oJK022 | Lambda red P1 | GTGTAGGCTGGAGCTGCTTC |
| oJK197 | lacZ-SSB-mYpet-KI_for | TGTGGAATTGTGAGCGGATAACAATTTCACACAGGAAACAGCT ATGGCCAGCAGAGGCGTAAA |
| oJK198 | lacZ-SSB-mYpet-KI_rev | TCATCATATTTAATCAGCGACTGATCCACCCAGTCCCAGACGA AGATGAATATCCTCCTTAGTTCCTA |
| oJK277 | rc_pKD4_9_30 | AATCGCTCAAGACGTGTAATGC |
| SO572 | pKD3/4-recG-KO-for | ACTGGTGGGCTACTATGCAGGCTGCAGGGTAAGTGCCATGGTG TAGGCTGGAGCTGCTT |
| SO573 | pKD3/4-recG-KO-rev | GTCTTCTCACTGCCGCCTTTTACGCATTCGAGTAACGTTCTAT GAATATCCTCCTTAGT |
| gbS282 | Gblock-pKD4-ssb-linker-Pamcherry | GCTTGCATGCAGATTGCAGCATTACACGTCTTGAGCGATTATG GCCAGCAGAGGCGTAAACAAGGTTATTCTCGTTGGTAATCTGG GTCAGGACCCGGAAGTACGCTACATGCCAAATGGTGGCGCAGT TGCCAACATTACGCTGGCTACTTCCGAATCCTGGCGTGATAAA GCGACCGGCGAGATGAAAGAACAGACTGAATGGCACCGCGTTG TGCTGTTCGGCAAACTGGCAGAAGTGGCGAGCGAATATCTGCG TAAAGGTTCTCAGGTTTATATCGAAGGTCAGCTGCGTACCCGT AAATGGACCGATCAATCCGGTCAGGATCGCTACACCACAGAAG TCGTGGTGAACGTTGGCGGCACCATGCAGATGCTGGGTGGTCG TCAGGGTGGTGGCGCTCCGGCAGGTGGCAATATCGGTGGTGGT CAGCCGCAGGGCGGTTGGGGTCAGCCTCAGCAGCCGCAGGGTG GCAATCAGTTCAGCGGCGGCGCGCAGTCTCGCCCGCAGCAGTC CGCTCCGGCAGCGCCGTCTAACGAGCCGCCGATGGACTTTGAT GATGACATTCCGTTCAGCTCGGCTGGCTCCGCTGCTGGTTCTG GCGAATTCATGGTTAGCAAGGGCGAGGAGGATAACATGGCCAT CATTAAGGAGTTCATGCGCTTCAAGGTGCACATGGAGGGGTCC GTGAACGGCCACGTGTTCGAGATCGAGGGCGAGGGCGAGGGCC GCCCCTACGAGGGCACCCAGACCGCCAAGCTGAAGGTGACCAA GGGTGGCCCCCTGCCCTTCACCTGGGACATCCTGTCCCCTCAA TTCATGTACGGCTCCAATGCCTACGTGAAGCACCCCGCCGACA TCCCCGACTACTTTAAGCTGTCCTTCCCCGAGGGCTTCAAGTG GGAGCGCGTGATGAAATTCGAGGACGGCGGCGTGGTGACCGTG ACCCAGGACTCCTCCCTGCAGGACGGTGAGTTCATCTACAAGG TGAAGCTGCGCGGCACCAACTTCCCCTCCGACGGCCCCGTAAT GCAGAAGAAGACCATGGGCTGGGAGGCCCTCTCCGAGCGGATG TACCCCGAGGACGGCGCCCTGAAGGGCGAGGTCAAGCCGAGAG TGAAGCTGAAGGACGGCGGCCACTACGACGCTGAGGTCAAGAC CACCTACAAGGCCAAGAAGCCCGTGCAGCTGCCCGGCGCCTAC |

| | | |
|---|---|---|
| | | AACGTCAACCGCAAGTTGGACATCACCTCACACAACGAGGACT ACACCATCGTGGAACAGTACGAACGTGCCGAGGGCCGCCACTC CACCGGCGGCATGGACGAGCTGTACAAGTAAGTGTAGGCTGGA GCTGCTTCGAAGTTCCTATACTTTCTAG |

### Table S11: Plasmids used in this study

| Designation | Containing Strain | Source or Reference |
|---|---|---|
| pKD3 | JEK42 | (1) |
| pKD4 | JEK45 | (1) |
| pSIM5 | JEK168 | (2) |
| pCP20 | JEK169 | (3) |
| pKD4-G4S-PAmCherry | JEK573 | (4) |
| pKD3-ssb-linker-PAmCherry | SE99 | This study |
| pSIM6 | S235 | (2) |

### Table S12: *Escherichia coli* bacterial strains used in this study

| Number | Designation or description | Relevant genotype | Construction or source strain designation | Reference |
|---|---|---|---|---|
| ET227 | Δ*lexA* Pol IV$^{T120P}$-PAmCherry SSB-mYPet | RW542 *dinB*$^{T120P}$-*PAmCherry-frt-kan-frt lacZ::ssb-mypet-frt* | Same | (5) |
| ET240 | Δ*lexA* SSB-PAmCherry | RW542 *lacZ::ssb-PAmCherry-frt-cat-frt* | P1*vir*: S146 → JEK418 | This study |
| ET340 | Δ*lexA* SSB-PAmCherry | RW542 *lacZ::ssb-PAmCherry-frt* | ET240 pCP20 Flp-FRT recombination | This study |
| ET350 | Δ*lexA* SSB-PAmCherry ε-mYPet | RW542 *lacZ::ssb-PAmCherry-frt dnaQ-mypet-frt-cat-frt* | P1*vir*: JEK466 → ET340 | This study |
| ET352 | Δ*lexA* Δ*dinB* | RW542 Δ*dinB::frt-kan-frt* | P1*vir*: JEK337 → JEK418 | This study |
| JEK5 | *E.coli* K-12 type strain | MG1655 | — | — |
| JEK168 | *E.coli* K-12 type strain with Cm$^{R}$ recombineering plasmid | pSIM5 in MG1655 | Same | (2, 4) |
| JEK170 | *E.coli* K-12 type strain with Amp$^{R}$ recombineering plasmid | pKD46 in MG1655 | Same | (1, 4) |
| JEK337 | Δ*dinB* | MG1655 Δ*dinB::frt-kan-frt* | Same | (4) |
| JEK418 | Δ*lexA* | *lexA51(Def) rpsL31 xyl-5 mtl-1 galK2 lacY1 tsx-33 supE44 thi-1 hisG4[Oc]* | RW542 | (6) |

| | | | | |
|---|---|---|---|---|
| | | *argE3[Oc] araD139 thr-1 Δ[gpt-proA]62 sulA211* | | |
| JEK466 | ε-mYPet | CH1358 *dnaQ-mypet-frt-cat-frt* | Same | (4) |
| JEK625 | Pol IV-PAmCherry | MG1655 *dinB-PAmCherry-frt-kan-frt* | Same | (4) |
| JEK726 | Pol IV$^{R,C}$-PAmCherry | MG1655 *dinB*$^{R,C}$*-PAmCherry-frt-kan-frt* | Same | (4) |
| JEK762 | Δ*lexA* SSB-mYPet | RW542 *lacZ::ssb-mypet-frt* | Same | (4) |
| JEK766 | Δ*lexA* Pol IV-PAmCherry SSB-mYPet | RW542 *dinB-PAmCherry-frt-kan-frt lacZ::ssb-mypet-frt* | Same | (4) |
| JEK790 | Δ*lexA* Pol IV$^{R,C}$-PAmCherry SSB-mYPet | RW542 *dinB*$^{R,C}$*-PAmCherry-frt-kan-frt lacZ::ssb-mypet-frt* | Same | (4) |
| S146 | SSB-PAmCherry | MG1655 *lacZ::ssb-PAmCherry-frt-cat-frt* | λ Red: SSB-PAmCherry-FRT-Cat-FRT → MG1655 pSIM5 | This study |
| S235 | *E.coli* K-12 type strain with Amp$^R$ recombineering plasmid | pSIM6 in MG1655 | Transformation: pSIM6 → JEK5 | (2) |
| S473 | Δ*recG* | MG1655 Δ*recG::frt-kan-frt* | λ Red: ΔrecG::FRT-Kan-FRT → MG1655 pSIM5 | This study |
| SCP004 | RecG-PAmCherry | MG1655 *recG-PAmCherry-frt-kan-frt* | λ Red: RecG-PAmCherry-FRT-Kan-FRT → MG1655 pSIM5 | This study |
| SCP007 | PriA-PAmCherry | MG1655 *priA-PAmCherry-frt-kan-frt* | λ Red: PriA-PAmCherry-FRT-Kan-FRT → MG1655 pSIM5 | This study |
| SCP008 | Δ*lexA* PriA-PAmCherry SSB-mYPet | RW542 *priA-PAmCherry-frt-kan-frt lacZ::ssb-mypet-frt* | P1*vir*: SCP007 → JEK762 | This study |
| SCP012 | Δ*lexA* RecG-PAmCherry SSB-mYPet | RW542 *recG-PAmCherry-frt-kan-frt lacZ::ssb-mypet-frt* | P1*vir*: SCP004 → JEK762 | This study |

### Supplementary Methods

*Detailed strain construction information*

**ET240/ET340:** Δ*lexA* SSB-PAmCherry. P1*vir* transduction was used to transfer the *ssb-PAmCherry* allele from strain S146 to strain JEK418. Plasmid pCP20 was then transformed into strain ET240 and Flp-FRT recombination was used to remove the Cm cassette, giving strain ET340.

**ET350:** Δ*lexA* SSB-PAmCherry ε-mYPet. P1*vir* transduction was used to transfer the *dnaQ-mYPet* allele from strain JEK466 to strain ET340.

**ET352:** Δ*lexA* Δ*dinB*. P1*vir* transduction was used to transfer the *ΔdinB::frt-kan-frt* allele from strain JEK337 to strain JEK418.

**S146:** MG1655 SSB-PAmCherry. In this construct, nucleotides nt1-1697 of *lacZ* were replaced with *ssb-PAmCherry-frt-cat-frt*. A backbone fragment of the plasmid pKD3 was amplified using the oligonucleotides Lambda red P1 and rc_pKD4_9_30 and combined with a Gene Block containing the SSAGSAAGSGEF linker and PAmCherry (Gblock-pKD4-ssb-linker-Pamcherry) by Gibson assembly, yielding plasmid pKD3-ssb-linker-PAmCherry. A recombineering fragment was then generated by amplifying this plasmid with primers lacZ-SSB-mYpet-KI_for and lacZ-SSB-mYpet-KI_rev and transformed into strain MG1655 pKD46 (JEK170), then pKD46 was cured to give strain S146.

**S473:** MG1655 Δ*recG*. In this construct, *recG* was replaced with *frt-kan-frt*, leaving only the first amino acid and the last 6 amino acids of *recG*. A recombineering fragment was generated by amplifying pKD4 with primers pKD3/4-recG-KO-for and pKD3/4-recG-KO-rev and transformed into S235 (MG1655 pSIM6), then pSIM6 was cured to give strain S473.

**SCP004:** MG1655 RecG-PAmCherry. A recombineering fragment was generated by amplifying pKD4-G4S-PAmCherry with primers recG-G4S-PAmC-KI-for and recG-PAmC-KI-rev and transformed into JEK168 (MG1655 pSIM5), then pSIM5 was cured to give strain SCP004.

**SCP007:** MG1655 PriA-PAmCherry. A recombineering fragment was generated by amplifying pKD4-G4S-PAmCherry with primers priA-G4S-PAmC-KI-for and priA-PAmC-KI-rev and transformed into JEK168 (MG1655 pSIM5), then pSIM5 was cured to give strain SCP007.

**SCP008:** Δ*lexA* PriA-PAmCherry SSB-mYPet. P1*vir* transduction was used to transfer the *priA-PAmCherry* allele from strain SCP007 to strain JEK762.

**SCP012:** Δ*lexA* RecG-PAmCherry SSB-mYPet. P1*vir* transduction was used to transfer the *recG-PAmCherry* allele from strain SCP004 to strain JEK762.
